# 5’-untranslated region sequences enhance plasmid-based protein production in *Sulfolobus acidocaldarius*

**DOI:** 10.1101/2024.03.01.582787

**Authors:** Laura Kuschmierz, Alexander Wagner, Tobias Busche, Jörn Kalinowski, Christopher Bräsen, Bettina Siebers

**Affiliations:** Molecular Enzyme Technology and Biochemistry (MEB), Environmental Microbiology and Biotechnology (EMB), Centre for Water and Environmental Research (CWE), Faculty of Chemistry, University of Duisburg-Essen, Essen, Germany; Microbial Genomics and Biotechnology, Center for Biotechnology (CeBiTec), Bielefeld University, Bielefeld, Germany

**Keywords:** protein expression, Archaea, *Sulfolobus acidocaldarius*, 5’-untranslated region, Shine-Dalgarno

## Abstract

*Sulfolobus acidocaldarius*, a thermoacidophilic archaeon of the phylum Thermoproteota (former Crenarchaeota), is a widely used model organism for gene deletion studies and recombinant protein production. Previous research has demonstrated the efficacy of the *saci_2122* promoter (P_ara_), providing low basal activity and high pentose-dependent induction. However, available expression vectors lack a 5’-terminal untranslated region (5’-UTR), which is a typical element in bacterial expression vectors, usually significantly enhancing protein production in bacteria. To establish *S. acidocaldarius* as a production strain in biotechnology in the long-term, it is intrinsically relevant to optimize its tools and capacities to increase production efficiencies. Here we show that protein production is increased by the integration of *S. acidocaldarius* 5’-UTRs into P_ara_ expression plasmids. Using the esterase Saci_1116 as a reporter protein, we observed a fourfold increase in soluble and active protein yield upon insertion of the *saci_1322* (*alba*) 5’-UTR. Screening of four additional 5’-UTRs from other highly abundant proteins (*thα*, *slaA*, *slaB, saci_0330*) revealed a consistent enhancement in target protein production. Additionally, site-directed mutagenesis of the Shine-Dalgarno (SD) motif within the *alba* 5’-UTR revealed its significance for protein synthesis. Ultimately, the *alba* 5’-UTR optimized expression vector demonstrated successful applicability in expressing various proteins, exemplified by its utilization for archaeal glycosyltransferases. Our results demonstrate that the integration of SD-motif containing 5’-UTRs significantly boosted plasmid-based protein production in *S. acidocaldarius*. This advancement in recombinant expression not only broadens the utility of *S. acidocaldarius* as an archaeal expression platform but also marks a significant step toward potential biotechnological applications.

## 1 Introduction

The efficient production of properly folded, functional proteins of interest (POIs) is fundamental in both basic research and biotechnology. A range of microorganisms and gene expression vectors are available for protein production. Most commonly, yeast or bacteria are used for gene overexpression, with *Escherichia coli* standing out as the most commonly used bacterial expression host (Rosano & Ceccarelli, 2014). Despite continuous advancements in expression systems, challenges such as suboptimal codon usage and posttranslational modifications, protein misfolding, and the formation of insoluble aggregates persist, hindering successful gene overexpression (de Lise et al., 2023).

Selecting an appropriate expression host is crucial for successful POI production. Besides eukaryotic or bacterial strains, Archaea offer high potential as protein production platforms. Archaea are characterized by a mosaic nature, sharing cellular features with both Bacteria and Eukaryotes, while additionally exhibiting unique archaeal properties. As Bacteria, Archaea are unicellular, lack organelles, and possess similar DNA structures (e.g. operon structures, one circular chromosome, plasmids). However, archaeal information processing (e.g. transcription and translation) and posttranslational modifications resemble simplified versions of eukaryotic processes (Lewis et al., 2021; Schocke et al., 2019; Esser et al., 2016; Jarrell et al., 2014, Eichler & Adams, 2005). While some hyperthermophilic proteins can be efficiently produced and purified using *Escherichia coli* with heat precipitation as an efficient purification step, the production of other thermophilic and/or archaeal proteins is difficult using classical bacterial expression hosts (e.g. Martinez-Espinosa, 2020; Straub et al., 2018). In these cases, proper expression or protein folding and stability may depend on special cofactors, posttranslational modifications, or suitable expression conditions, such as temperature (Eichler & Adams, 2005; Martinez-Espinosa, 2020; Straub et al., 2018). Thus, expression in the native or in a closely related organism was suggested as alternative for efficient protein synthesis (Straub et al., 2018).

With its established genetic system and genomic stability, the thermoacidophilic, aerobic archaeon *Sulfolobus acidocaldarius* (T_opt_ 70 - 75°C, pH_opt_ 2 - 3; Brock et al., 1972; Chen et al., 2005) offers a versatile platform for gene deletion studies and overexpression (de Lise et al., 2023, Wagner et al., 2012; Lewis et al., 2021). Inducible expression vectors utilizing sugar-inducible promoters (e.g. P_ara_ (L-arabinose) from *saci_2122*, P_xyl_ (D-xylose) from *saci_1938* or P_mal_ (maltose) from *saci_1165*) have been developed, featuring various N- or C-terminal protein tags (see van der Kolk et al., 2020 for an overview) for successful protein expression (e.g. Stracke et al., 2020; Zweerink et al., 2017). In a comparative promoter screening, P_ara_ was identified as the most efficient promoter, exhibiting the highest pentose-dependent induction with D-xylose and L-/D-arabinose and low basal activity (van der Kolk et al., 2020). While D-xylose and L-arabinose are metabolized by *S. acidocaldarius*, D-arabinose does not serve as a carbon source and can therefore act as an artificial inducer.

In general, mRNAs can carry UTRs at the 5’ and 3’-end. Both are known to be involved in many important regulatory processes, including translation initiation, transcript stability, mRNA secondary structure formation, and posttranscriptional regulation, e.g. through the action of RNA binding proteins or small regulatory RNAs (Brenneis & Soppa, 2009; Gomes-Filho et al., 2018; Ren et al., 2017). Notably, under the control of the native P_ara_ sequence, *S. acidocaldarius* generates leaderless gene transcripts, lacking a 5’-untranslated region (UTR) (Cohen et al., 2016). By definition, leaderless mRNAs may either directly start with the initiation codon or possess up to five (Babski et al., 2016) or ten (Brenneis et al., 2007; Chen et al., 2015) 5’-terminal nucleotides upstream of the translation start codon (Fig. 1). In contrast, leadered mRNAs carry a 5′-UTR, i.e. a non-coding regulatory sequence at the 5’-end of the transcript. 5’-UTRs may contain a Shine-Dalgarno (SD) sequence motif (Hering et al., 2009; Srivastava et al., 2016; Wurtzel et al., 2010). If present, the SD sequence (core sequence: GGAGGU (Shine & Dalgarno, 1975)) enables base pairing with the anti-Shine-Dalgarno sequence (aSD) at the 3’-end of the 16S rRNA of the small ribosomal subunit (Ma et al., 2002; Torarinsson et al., 2005), thereby supporting translation initiation. Based on the direct interaction of SD to aSD, the SD is referred to as ribosome binding site (RBS). However, in the process of translation initiation, the ribosome covers an mRNA region of approx. 30 nts, defined as ribosome docking site (RDS). This docking area includes the SD sequence, the translation start codon, an intermediate spacer region (of variable length and nucleotide composition), and the first nucleotides of the coding sequence (Reeve et al., 2014; Chen et al., 1994) (Fig. 1).

**Figure 1:**
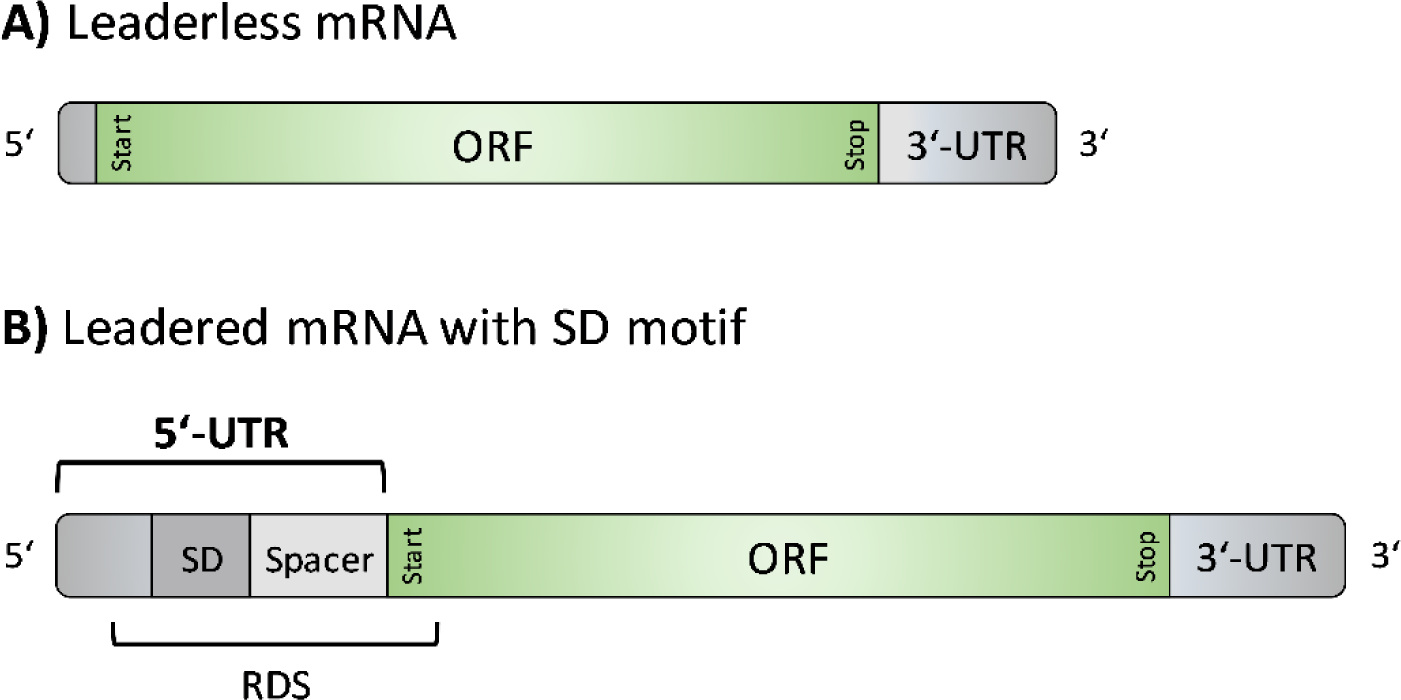
Schematic view of structural elements of leaderless mRNAs and leadered mRNAs with a Shine-Dalgarno motif. **A**) Leaderless mRNAs are composed of an open reading frame (ORF), defined by a translation start and stop codon, and a 3’-terminal untranslated region (UTR). Optionally, up to ten nucleotides may be localized at the 5’ end (Brenneis et al., 2007). **B**) Leadered mRNAs feature a 5’-terminal UTR (more than ten 5’-terminal nucleotides), which may contain a Shine-Dalgarno (SD) sequence that allows direct interaction with the aSD sequence of the 16S rRNA. The region between SD and translation start codon is defined as spacer region. The ribosome docking site (RDS) includes SD, spacer and the first nucleotides of the coding region (Reeve et al., 2014; Chen et al., 1994).

The use of a 5’-UTR, including a preserved SD sequence, is common for many bacterial expression vectors, and in Bacteria the insertion, enlargement, or modification of the 5’-UTR (RBS) has been shown to increase plasmid-encoded protein production significantly (e.g. Liebeton et al., 2014; Vellanoweth & Rabinowitz, 1992; Volkenborn et al., 2020). In contrast, their utilization in archaeal expression plasmids remains less explored (e.g. Brenneis et al., 2007; Akinyemi et al., 2021). In this study, we systematically analyze the effect of 5’-UTR insertions into *S. acidocaldarius* expression plasmids for the production of two reporter proteins, esterase Saci_1116 and β-galactosidase LacS from *Saccharolobus solfataricus*. Additionally, we employ 5’-UTR-optimized expression vectors with N- or C-terminal Twin-Strep tags for the production of archaeal glycosyltransferases.

## 2 Materials and Methods

### Strains and growth conditions

The uracil auxotrophic mutant *S. acidocaldarius* MW001 (Wagner et al., 2012) was used as the expression host. The strain originates from *S. acidocaldarius* DSM639, but lacks 322 bp of *pyrE* (*saci_1597*), enabling genetic selection. Cultures were grown aerobically in Brock medium (Brock et al., 1972) at pH 3.0, 140 rpm, and 76°C (New Brunswick Innova 44 incubator, Eppendorf, Germany). Cell growth was monitored by turbidity measurements at 600 nm. *S. acidocaldarius* MW001 cultures were supplemented with 10 µg/mL uracil (min. 99%, Merck, Germany). Transformed clones of *S. acidocaldarius* MW001, carrying an expression plasmid with *pyrEF* from *Saccharolobus solfataricus*, were grown without the addition of uracil.

*Escherichia coli* DH5α and ER1821 (New England Biolabs (NEB), USA) were used for cloning and methylation of *S. acidocaldarius* expression plasmids, respectively. The strains were grown in Lysogeny broth (LB) medium (Carl Roth, Germany), supplemented with respective antibiotics, at 37°C (and 180 rpm for liquid cultures).

### Plasmid construction

pBSaraFX-xxxUTR-Nt/Ct-SS plasmids are originally based on pSVAaraFX-Nt/Ct-S S expression vectors (van der Kolk et al., 2020). They possess the vector backbone of *Sulfolobus islandicus* plasmid pRN1 (Lipps, 2004) and contain basal shuttle vector elements such as origins of replication for *S. acidocaldarius* and *E. coli*, as well as *pyrEF* from *S. solfataricus* and an ampicillin resistance gene (β-lactamase) for selection in *S. acidocaldarius* or *E. coli*, respectively. The gene of interest expression cassette is flanked by different restriction sites and is composed of the pentose-inducible promoter of *saci_2122* (P_ara_) and *lacI*, *lacZ* as a reporter system enabling blue/white screening in *E. coli*. The vector encodes a Twin-Strep tag, enabling N- or C-terminal protein tagging (pSVAaraFX-NtSS or - CtSS; van der Kolk et al., 2020). “FX” indicates that the FX cloning strategy can be applied (class IIS restriction enzyme *Sap*I; Geertsma & Dutzler, 2011).

### Integration of different 5’-UTR sequences into *saci_1116* expression plasmid

In this study, the pSVAaraFX-*saci_1116*-CtSS expression plasmid was expanded by the insertion of different 5’-untranslated region sequences that were integrated directly upstream of the translation start codon of the reporter gene *saci_1116* (Fig. S1, S2). The insertion of a 5’-UTR was achieved using standard cloning procedures: modified promoter-5’-UT R fragments were PCR amplified with a fwd-primer (UTR-fwd, Tab. S1), that bound to the vector backbone upstream of the *saci_2122* promoter, in combination with a rev-primer containing the desired 5’-UTR sequence and a *Nco*I restriction site that bound upstream of the translation start codon (Tab. S1). pSVAaraFX-CtSS served as PCR template. The resulting PCR product consisted of a fragment of the vector backbone, carrying a *Sac*II restriction site, P_ara_, and the newly integrated 5’-UTR sequence with a *Nco*I restriction site. The fragment was cloned into pSVAaraFX-*saci_1116*-CtSS by restriction with *Sac*II and *Nco*I (NEB) followed by ligation (T4 DNA ligase, NEB). The resulting *saci_1116* expression plasmid, containing a 5’-UTR, was named pBSaraFX-xxxUTR-*saci_1116*-CtSS. Successful cloning was confirmed by sequencing (LGC genomics, Germany).

### Generation of *lacS* expression constructs with different 5’-UTR sequences

The series of expression plasmids for *saci_1116*, each containing distinct 5’-UTR sequences, served as the basis for the development of corresponding *lacS* expression plasmids. Amplification of *lacS* from *Saccharolobus solfataricus* (SSO3019) was conducted using primers *lacS*-SSO-*Nco*I-fwd and *lacS*-SSO-*Xho*I-rev (Tab. S1). Subsequently, the resulting PCR product and the pBSaraFX-xxxUTR-*saci_1116*-CtSS expression plasmids (Tab. 1) underwent restriction enzyme digestion with *Nco*I and *Xho*I. This process facilitated the excision of *saci_1116* from the xxxUTR-plasmid backbone, excluding it from subsequent cloning steps through agarose gel extraction of the plasmid fragment. The *Nco*I-*Xho*I digested pBSaraFX-xxxUTR-CtSS fragment and *lacS* were ligated. Verification of the resultant plasmid set, designated as pBSaraFX-xxxUTR-*lacS*-CtSS, was conducted through sequencing. Subsequently, this plasmid set was utilized to investigate LacS production in *S. acidocaldarius*.

**Table 1:**
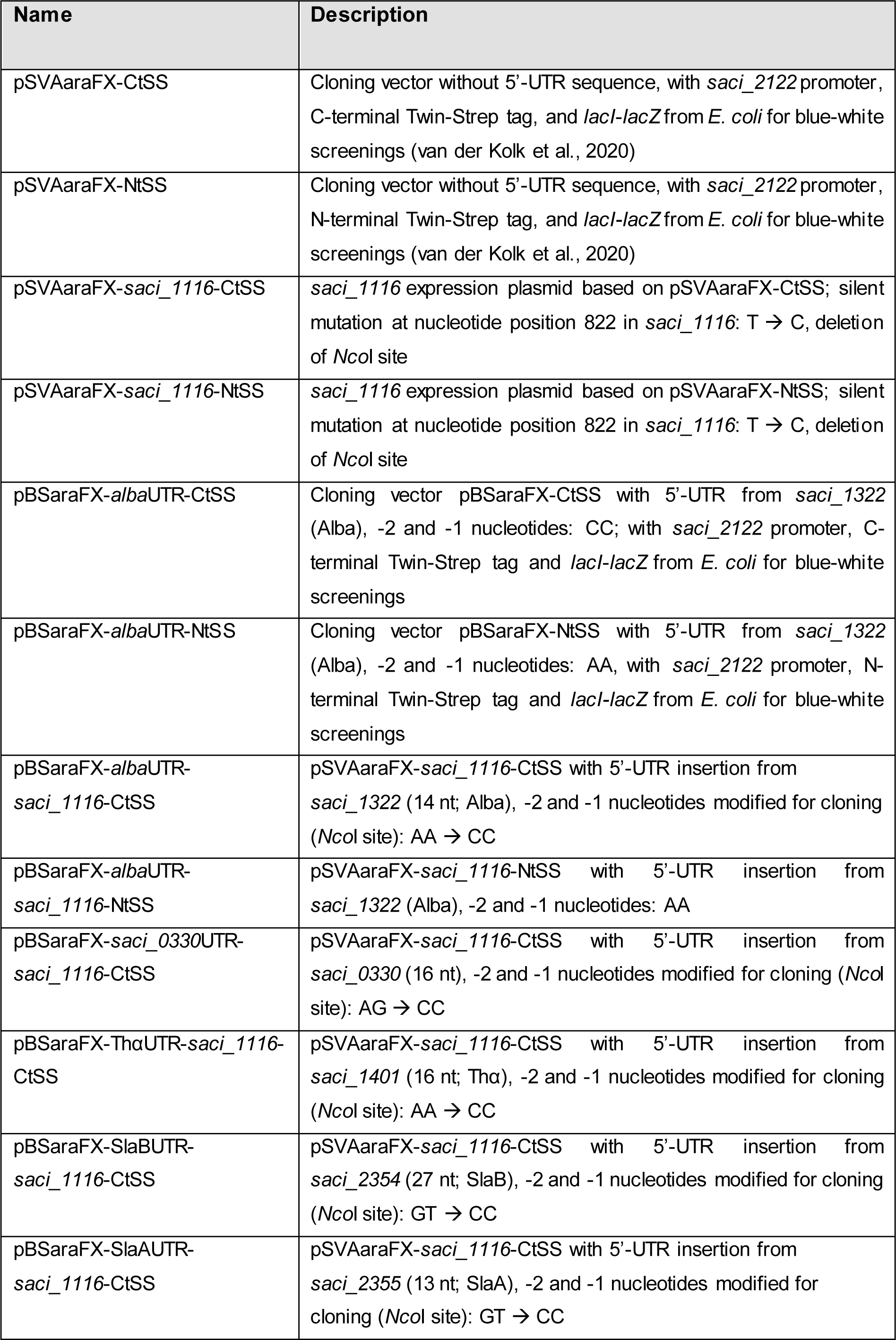
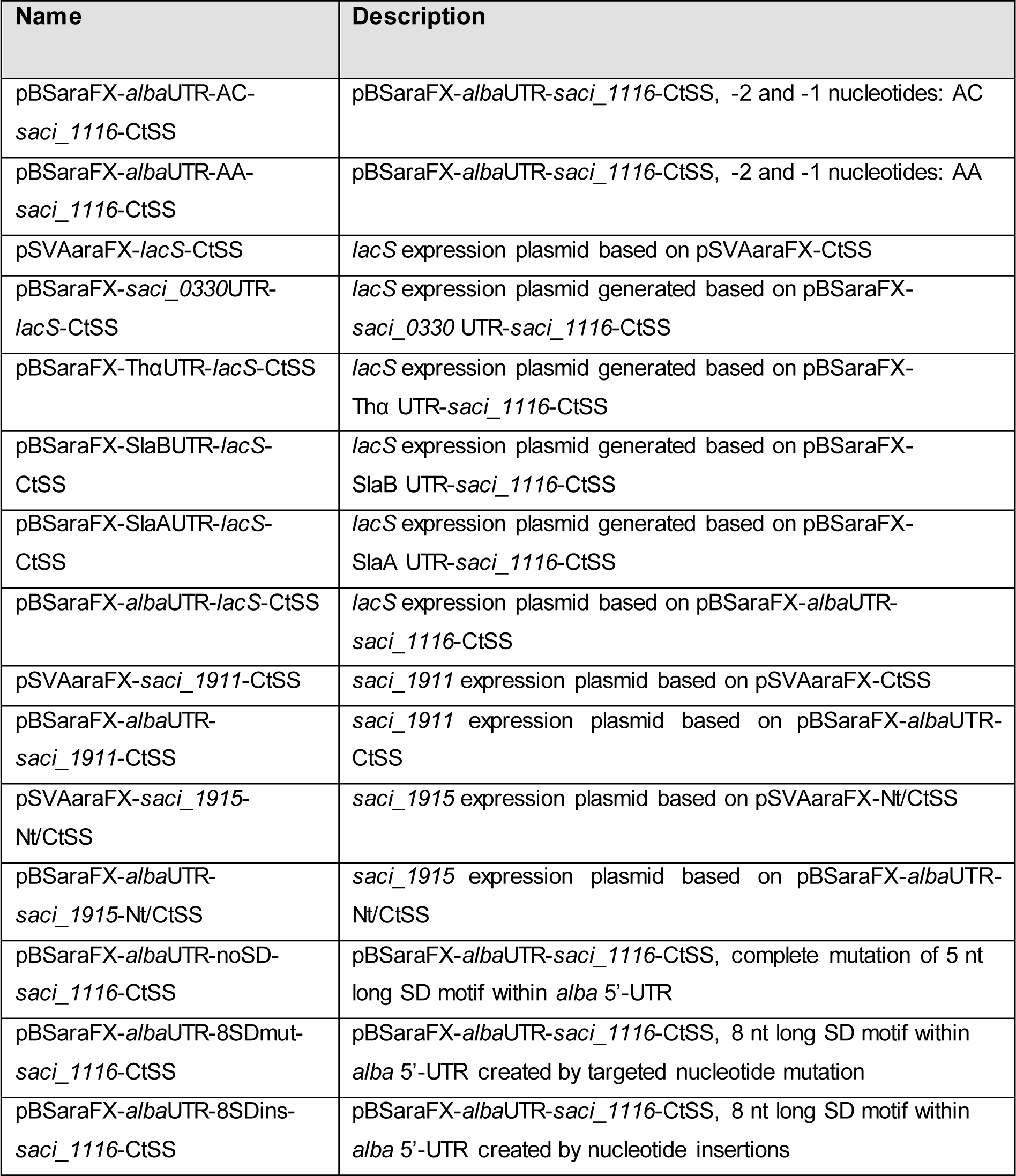
Plasmids used in this study.

| Name | Description |
| --- | --- |
| pSVAaraFX-CtSS | Cloning vector without 5'-UTR sequence, with *saci_2122* promoter, C-terminal Twin-Strep tag, and *lacI-lacZ* from *E. coli* for blue-white screenings (van der Kolk et al., 2020) |
| pSVAaraFX-NtSS | Cloning vector without 5'-UTR sequence, with *saci_2122* promoter, N-terminal Twin-Strep tag, and *lacI-lacZ* from *E. coli* for blue-white screenings (van der Kolk et al., 2020) |
| pSVAaraFX-*saci_1116*-CtSS | *saci_1116* expression plasmid based on pSVAaraFX-CtSS; silent mutation at nucleotide position 822 in *saci_1116*: T → C, deletion of *NcoI* site |
| pSVAaraFX-*saci_1116*-NtSS | *saci_1116* expression plasmid based on pSVAaraFX-NtSS; silent mutation at nucleotide position 822 in *saci_1116*: T → C, deletion of *NcoI* site |
| pBSaraFX-*alba*UTR-CtSS | Cloning vector pBSaraFX-CtSS with 5'-UTR from *saci_1322* (Alba), -2 and -1 nucleotides: CC; with *saci_2122* promoter, C-terminal Twin-Strep tag and *lacI-lacZ* from *E. coli* for blue-white screenings |
| pBSaraFX-*alba*UTR-NtSS | Cloning vector pBSaraFX-NtSS with 5'-UTR from *saci_1322* (Alba), -2 and -1 nucleotides: AA, with *saci_2122* promoter, N-terminal Twin-Strep tag and *lacI-lacZ* from *E. coli* for blue-white screenings |
| pBSaraFX-*alba*UTR-*saci_1116*-CtSS | pSVAaraFX-*saci_1116*-CtSS with 5'-UTR insertion from *saci_1322* (14 nt; Alba), -2 and -1 nucleotides modified for cloning (*NcoI* site): AA → CC |
| pBSaraFX-*alba*UTR-*saci_1116*-NtSS | pSVAaraFX-*saci_1116*-NtSS with 5'-UTR insertion from *saci_1322* (Alba), -2 and -1 nucleotides: AA |
| pBSaraFX-*saci_0330*UTR-*saci_1116*-CtSS | pSVAaraFX-*saci_1116*-CtSS with 5'-UTR insertion from *saci_0330* (16 nt), -2 and -1 nucleotides modified for cloning (*NcoI* site): AG → CC |
| pBSaraFX-ThaUTR-*saci_1116*-CtSS | pSVAaraFX-*saci_1116*-CtSS with 5'-UTR insertion from *saci_1401* (16 nt; Tha), -2 and -1 nucleotides modified for cloning (*NcoI* site): AA → CC |
| pBSaraFX-SlaBUTR-*saci_1116*-CtSS | pSVAaraFX-*saci_1116*-CtSS with 5'-UTR insertion from *saci_2354* (27 nt; SlaB), -2 and -1 nucleotides modified for cloning (*NcoI* site): GT → CC |
| pBSaraFX-SlaAUTR-*saci_1116*-CtSS | pSVAaraFX-*saci_1116*-CtSS with 5'-UTR insertion from *saci_2355* (13 nt; SlaA), -2 and -1 nucleotides modified for cloning (*NcoI* site): GT → CC |
| pBSaraFX-*alba*UTR-AC-*saci_1116*-CtSS | pBSaraFX-*alba*UTR-*saci_1116*-CtSS, -2 and -1 nucleotides: AC |
| pBSaraFX-*alba*UTR-AA-*saci_1116*-CtSS | pBSaraFX-*alba*UTR-*saci_1116*-CtSS, -2 and -1 nucleotides: AA |
| pSVAaraFX-*lacS*-CtSS | *lacS* expression plasmid based on pSVAaraFX-CtSS |
| pBSaraFX-*saci_0330*UTR-*lacS*-CtSS | *lacS* expression plasmid generated based on pBSaraFX-*saci_0330* UTR-*saci_1116*-CtSS |
| pBSaraFX-ThaUTR-*lacS*-CtSS | *lacS* expression plasmid generated based on pBSaraFX-Tha UTR-*saci_1116*-CtSS |
| pBSaraFX-SlaBUTR-*lacS*-CtSS | *lacS* expression plasmid generated based on pBSaraFX-SlaB UTR-*saci_1116*-CtSS |
| pBSaraFX-SlaAUTR-*lacS*-CtSS | *lacS* expression plasmid generated based on pBSaraFX-SlaA UTR-*saci_1116*-CtSS |
| pBSaraFX-*alba*UTR-*lacS*-CtSS | *lacS* expression plasmid based on pBSaraFX-*alba*UTR-*saci_1116*-CtSS |
| pSVAaraFX-*saci_1911*-CtSS | *saci_1911* expression plasmid based on pSVAaraFX-CtSS |
| pBSaraFX-*alba*UTR-*saci_1911*-CtSS | *saci_1911* expression plasmid based on pBSaraFX-*alba*UTR-CtSS |
| pSVAaraFX-*saci_1915*-Nt/CtSS | *saci_1915* expression plasmid based on pSVAaraFX-Nt/CtSS |
| pBSaraFX-*alba*UTR-*saci_1915*-Nt/CtSS | *saci_1915* expression plasmid based on pBSaraFX-*alba*UTR-Nt/CtSS |
| pBSaraFX-*alba*UTR-noSD-*saci_1116*-CtSS | pBSaraFX-*alba*UTR-*saci_1116*-CtSS, complete mutation of 5 nt long SD motif within *alba* 5'-UTR |
| pBSaraFX-*alba*UTR-8SDmut-*saci_1116*-CtSS | pBSaraFX-*alba*UTR-*saci_1116*-CtSS, 8 nt long SD motif within *alba* 5'-UTR created by targeted nucleotide mutation |
| pBSaraFX-*alba*UTR-8SDins-*saci_1116*-CtSS | pBSaraFX-*alba*UTR-*saci_1116*-CtSS, 8 nt long SD motif within *alba* 5'-UTR created by nucleotide insertions |

### Site-directed mutagenesis of *alba* 5’-UTR

To assess the impact of −2 and −1 nucleotides, located directly upstream of the translation start codon within the *alba* 5’-UTR, on gene expression, we employed site-directed mutagenesis. Plasmid pBSaraFX-*alba*UTR-*saci_1116*-CtSS served as the template for modifications (Tab. 1). Each primer contained the desired mutation and was complementary to the other (Tab. S1). To prevent the formation of primer homodimers in the reaction mixture, DNA amplification was performed in single-primer reactions, following the protocol outlined by Edelheit et al., 2009. Q5^®^ High-Fidelity DNA Polymerase and its recommended reaction buffer (NEB) were utilized for the amplification process.

Correspondingly, the SD sequence motif within the *alba* 5’-UTR was modified by site-directed mutagenesis, as described by Edelheit et al., 2009 (Tab. S1) using pBSaraFX-*alba*UTR-*saci_1116*-CtSS as template (“noSD” and “8SDmut” variants of *alba* 5’-UTR (Tab. 1; Fig. 6A)). The “8SDins” variant, designated as pBSaraFX-*alba*UTR-8SDins-*saci_1116*-CtSS, was constructed via restriction and ligation-dependent cloning. Initially, PCR was conducted using primers SD-fwd and 8SD-ins-rev with pSVAaraFX-CtSS serving as the template (Tab. S1, Tab. 1). Subsequently, the resulting PCR product and the vector pSVAaraFX-*saci_1116*-CtS S underwent restriction using *Sac*II and *Nco*I, followed by ligation.

### Generation of cloning vectors pBSaraFX-*alba*UTR-CtSS and -NtSS

Generally, *S. acidocaldarius* expression vectors contain *lacI* and *lacZ* in the expression cassette for blue/white screening in *E. coli*. To enable this feature to be used in the cloning process of *alba* 5’-UTR containing plasmids, two *alba* 5’-UTR cloning vectors with inserted *lacI*, *lacZ*, and an N- or C-terminal Twin-Strep tag were generated. pBSaraFX-*alba*UTR-NtSS (N-terminal Twin-Strep tag) was generated by the insertion of the original *alba* (*saci_1322*) 5’-UTR (5’ GATAGGTGGTTTAA 3’) directly upstream of the gene expression cassette start codon into pSVAaraFX-NtSS by overlap PCR using the primer pair *saci_1322*-UTR-NtSS-fwd and -rev (Tab. S1). To obtain the cloning vector pBSaraFX-*alba*UTR-CtSS (C-terminal Twin-Strep tag), the *saci_1116* expression plasmid pBSaraFX-*alba*UTR-*saci_1116*-CtSS as well as the vector pSVAaraFX-CtSS, containing *lacI* and *lacZ,* were restricted with *Nco*I and *Xho*I. The obtained vector backbone and the *lacI-lacZ* fragment were ligated, resulting in cloning vector pBSaraFX-*alba*UTR-CtSS (containing the modified *alba* 5’-UTR: 5’ GATAGGTGGTTT<u>CC</u> 3’).

### Cloning of gene expression plasmids for glycosyltransferases

Genes of interest, namely *saci_1911* and *saci_1915* from *S. acidocaldarius* were cloned into pBSaraFX-*alba*UTR-Nt/CtSS expression vectors (Tab. 1). All cloning primers and the used restriction sites are included in Tab. S1.

### Transformation of *S. acidocaldarius* MW001

Competent cells were prepared according to Wagner et al., 2012, and transformed with 300-500 ng of methylated plasmid DNA using a Gene Pulser Xcell (Bio-Rad, Germany) at 2000 V, 600 Ω and 25 µF in 1 mm cuvettes. Cell recovery was performed in Brock medium (pH 5.0), supplemented with 0.1% (*w/v*) N-Z-amine (casein enzymatic hydrolysate N-Z-Amine® AS, Merck, Germany), for 30 min-4 h at 76°C in a heat block (speed of 300 rpm; ThermoMixer F1.5, Eppendorf, Germany) (Wagner et al., 2014). Cells were plated on Gelrite-Brock plates with 0.6% (*w/v*) Gelrite® (Gellan Gum: K9A-40, Serva Electrophoresis, Heidelberg, Germany), 0.1% (*w/v*) N-Z-amine and 0.2% (*w/v*) dextrin (pure, Carl Roth, Germany), lacking uracil. Plates were incubated in a closed metal box for 5-6 d at 76°C.

### *S. acidocaldarius* MW001 expression cultures

After transformation of *S. acidocaldarius* MW001 and incubation of cells on Gelrite-Brock plates, clones were transferred to liquid Brock medium (0.1% (*w/v*) N-Z-amine (Merck, Germany) and 0.2% (*w/v*) dextrin (Carl Roth, Germany), pH 3.0), and cultures were grown to the late log growth phase (OD_600nm_ 0.9-1.2) at 140 rpm and 76°C. The presence of the expression plasmid was verified by colony PCR using gene-specific primer pairs (Tab. S1). Positive cell cultures, carrying an expression plasmid with SSO*pyrEF*, were used to inoculate pre-cultures for *S. acidocaldarius* expression. Pre-cultures were cultivated in Brock medium supplemented with 0.1% (*w/v*) N-Z-amine (Merck, Germany) and 0.2% (*w/v*) dextrin (Carl Roth, Germany) without uracil until the log growth phase. Expression cultures were inoculated to an OD_600nm_ of 0.05-0.1, and 0.1% (*w/v*) N-Z-amine as well as 0.2% or 0.3% (*w/v*) D-xylose (min. 99%, Carl Roth, Germany) was added to the medium. Expression cultures were grown at 140 rpm and 76°C (New Brunswick Innova 44 incubator, Eppendorf, Germany). Cells were harvested by centrifugation (7000 *x* g, 4°C, 20 min) at OD_600nm_ values of 0.6 or 0.9-1.2, as stated in the results section. Cell pellets were stored at −80°C until further use.

### Sodium dodecyl sulfate-polyacrylamide gel electrophoresis (SDS-PAGE) and immunoblotting of *S. acidocaldarius* cell lysate samples

*S. acidocaldarius* cell samples were adjusted to a theoretical OD_600nm_ of 10 by resuspension of cells in appropriate volumes of 50 mM TRIS/HCl pH 7.5 with 0.5% (v/v) Triton X-100 (Gerbu, Germany). 5x SDS-PAGE loading dye (125 mM TRIS/HCl pH 6.8, 4% (w/v) SDS, 20% (v/v) glycerol, 10% (v/v) β-mercaptoethanol, 0.01% (w/v) Bromophenol Blue) was added and samples were heated for 5 min at 99 °C. 5 µl of PageRuler Unstained or PageRuler (Plus) Prestained Protein Ladder (Thermo Fisher Scientific, USA) and 20 µl of each cell lysate sample (approx. 30 µg of protein) were applied to SDS-PAGE. Gels were either stained by Coomassie Brilliant Blue staining (0.05% (*w/v*) Coomassie Brilliant Blue G-250 (Merck, Germany), 40% (*v/v*) ethanol, 10% (*v/v*) acetic acid) and imaged using the Molecular Imager Gel Doc XR System and the Quantity One Software Package (Bio-Rad, Germany), or they were used for immunodetection. Proteins were transferred on a PVDF membrane (Roti-PVDF, pore size 0.45 µm, Carl Roth, Germany) using blotting buffer (50 mM TRIS, 40 mM glycine, 0.1% (*w/v*) SDS, 20% (*v/v*) EtOH (p.a.)) and the Bio-Rad trans-blot® TurboTM Trans System (25 V, 0.5-1 A, 25 min; Bio-Rad, Germany). Afterward, the membrane was blocked with 0.2% (*w/v*) Tropix^®^ I-BLOCK (Applied Biosystems, Thermo Fisher Scientific, USA) in TBS-T buffer (20 mM TRIS-HCl pH 7.5, 500 mM NaCl, 0.05% (*v/v*) Tween 20) at 4°C overnight. The next day, the membranes were floated in a solution of 1:50000 Strep-Tactin HRP conjugate (IBA Lifesciences, Germany) with 0.1% (*w/v*) I-BLOCK in TBS-T buffer and incubated while shaking at RT for 1-2 h. The membrane was washed twice using 0.1% (*w/v*) I-BLOCK in TBS-T buffer for 10 min each, followed by two washing steps using TBS-T and TBS buffer, respectively. Clarity Western ECL Substrate (Bio-Rad, Germany) and the VersaDoc MP-4000 imaging system (Bio-Rad, Germany) were used for immunodetection and documentation.

### Bicinchoninic acid (BCA) protein assay of *S. acidocaldarius* cell lysates

*S. acidocaldarius* cell suspensions were adjusted to a theoretical OD_600nm_ of 5 by resuspension of cells in 50 mM TRIS/HCl pH 7.5 with 0.5% (v/v) Triton X-100 (Gerbu, Germany). Cells were lysed by sonication (ultrasonic processor UP 200s (Hielscher Ultrasonics, Germany)) with a cycle of 0.5 s^-1^ and an amplitude of 50% for 1 min on ice. Protein concentrations in cell lysate samples were determined using the bicinchoninic acid (BCA) assay (Uptima BC assay protein quantification kit, France). Cell lysates (25 µL) and bovine serum albumin (BSA) protein standards (0-1 mg/mL BSA (Merck, Germany) in resuspension buffer) were transferred in the wells of a 96-well microtiter plate (MTP; flat bottom, polystyrene, Sarstedt, Germany) and mixed with 200 µL BCA solution each. For each biological replicate three technical replicates were analyzed. After 3 h of incubation at room temperature absorption at 562 nm was determined in a Tecan plate reader (Tecan Infinite^®^ M200, NanoQuant PlateTM; Tecan, Switzerland).

### Esterase activity assay (Saci_1116)

To determine the esterase activity of Saci_1116 in cell lysates of *S. acidocaldarius* cultures, *para*-nitrophenyl acetate (*p*NPA, Merck, Germany) was used as a substrate (Sobek & Görisch, 1988 and 1989). 175 µL of 50 mM TRIS/HCl (pH 7.5) and 5 µL of 10-fold diluted *S. acidocaldarius* cell lysate sample (diluted with reaction buffer) were pipetted into the wells of a 96-well MTP (flat bottom, polystyrene, Sarstedt, Germany) and pre-warmed in the Tecan plate reader at 42°C for 10 min (Tecan Infinite^®^ M200, NanoQuant PlateTM; Tecan, Switzerland). Afterward, 20 µl of 10 mM *p*NPA (dissolved in acetonitrile) were added to the wells by automatic dispersion, yielding a substrate concentration of 1 mM *p*NPA in the assay. Measurements were performed in 60 s cycles at 410 nm for 1 h, following the formation of *p*NP (4-nitrophenol). *p*NP (Merck, Germany) calibration curves were conducted under the same conditions. Experiments were performed in three biological and technical replicates. Enzyme activity measurements of purified esterase Saci_1116-CtSS were performed under the same assay conditions, using approx. 40 and 20 ng of pure protein. The initial reaction velocities were used for the calculation of activities. *p*NP production was calculated using the established *p*NP calibration curve. Specific enzyme activities (U/mg of protein) were determined by the protein concentration obtained in the BCA assay. Student’s t-test was performed to determine significant differences between the measurements of specific activity under the influence of different 5’-UTR sequences (absence or presence of *alba* 5’-UTR, *alba* 5’-UTR with different - 2 and −1 nucleotide compositions, use of different 5’-UTR sequences). In general, the null hypothesis was: the esterase activity (protein production) is the same in both compared expression conditions. The null hypothesis was rejected at a 95% confidence level if the p-value was less than 0.05, indicating that there was a statistically significant difference between two conditions.

### β-Galactosidase activity assay (LacS)

Cells of *S. acidocaldarius* expression cultures were resuspended in 50 mM sodium phosphate buffer (pH 6.5) supplemented with 50 mM NaCl and disrupted using prefilled 0.1 mm Glass Bead tubes and the Precellys 24 tissue homogenizer (Bertin technologies, France). The cell lysis process involved three cycles of homogenization at 6.500 *x*g for 20 s each, with intermittent cooling on ice for 1 min between cycles. Subsequently, cell lysates were centrifuged at 21428 *x*g at 4°C for 40 min to obtain the supernatant, the crude extract, which was next utilized for protein concentration determination. Protein concentration was assessed using the Bradford protein assay (QuickStart^TM^, Bio-Rad, Germany) with BSA (Sigma-Aldrich, USA) as protein standard (Zor & Selinger, 1996). LacS activity in crude extracts was determined using a continuous assay in the BioTek Synergy H1 microplate reader with 96-well microtiter plates (flat bottom, polystyrene, Thermo Fisher Scientific, USA). The wells of a 96-well MTP were filled with 177.5 µL of 50 mM sodium phosphate buffer (pH 6.5) and 2.5 µL of crude extract and pre-warmed in the reader at 70°C for 20 min. Then, 20 µl of 10 mM *para-* nitrophenyl β-D-galactopyranoside (dissolved in buffer) were automatically dispensed into the wells, resulting in a final substrate concentration of 1 mM *p*NPG in the assay. The plate was shaken for 5 s, and measurements were performed in 60 s cycles at 410 nm for 30 min to monitor the formation of *p*NP (4-nitrophenol). Calibration curves for *p*NP were prepared under the same conditions. Experimental procedures were replicated in three biological and technical replicates. Finally, the initial linear range of *p*NP formation following the enzymatic reaction (8 min) was utilized to calculate specific enzyme activity.

### Purification of Twin-Strep-tagged proteins from *S. acidocaldarius* crude extracts

Purification was performed according to the manufacturer’s instructions (IBA Lifesciences, Germany). Briefly, *S. acidocaldarius* cell suspensions were adjusted to a theoretical OD_600nm_ of 5 by resuspension in buffer W (100 mM TRIS/HCl pH 8.0, 150 mM NaCl). Cells were disrupted by sonication (UP 200s, Hielscher Ultrasonics, Germany) with a cycle of 0.5 s^-1^ and an amplitude of 55% three times for 5 min each on ice (cooling pauses of 1 min each) and cell lysates were centrifuged at 16,000 *x* g and 4°C for 40 min to remove cell debris. Supernatants were filtered (0.45 µm polyvinylidene fluoride membrane, Carl Roth, Germany) and Twin-Strep tagged proteins of interest were purified from crude extracts (soluble fraction) using the Strep-Tactin^®^XT Superflow^®^ Kit (IBA Lifesciences, Germany). Purification was performed according to the manufacturer’s instructions, using buffer BXT (100 mM TRIS/HCl pH 8.0, 150 mM NaCl, 1 mM EDTA, 50 mM biotin) as elution buffer. Protein purity was analyzed by SDS-PAGE and the protein concentration was determined using the Bradford assay (QuickStart^TM^, Bio-Rad, Germany) with BSA as the protein standard (Zor & Selinger, 1996).

## 3 Results

### Influence of the *alba* 5’-UTR on the plasmid-encoded production of esterase Saci_1116

In previous studies, promoter P_ara_ (*saci_2122*) has demonstrated its efficacy in providing protein production in *S. acidocaldarius*, characterized by high pentose-dependent induction and low basal activity (van der Kolk et al.; 2020). Notably, RNA sequencing data (Cohen et al., 2016; https://exploration.weizmann.ac.il/TCOL/) indicate that this promoter sequence leads to the generation of leaderless gene transcripts. This prompted our investigation into whether the incorporation of a 5’-UTR (RBS) could enhance plasmid-based protein production. To assess the impact of a 5’-UTR on protein yield, we utilized the 5’-UTR sequence of the Alba encoding gene *saci_1322* (*alba*) from *S. acidocaldarius*. Alba (Acetylation Lowers Binding Affinity) (Laurens et al., 2012) is a well-conserved archaeal double-stranded DNA/RNA binding protein (Wardleworth et al., 2002; Cajili & Prieto, 2022) and it was already reported to be highly abundant in *Sulfolobus* (4.8% of cellular protein in *Sulfolobus shibatae* (Mai et al., 1998)). In agreement, available full proteome data from *S. acidocaldarius* identified it as the most abundant cellular protein under different growth conditions (label-free quantification intensity, 0.2% (w/v) NZA and 0.2% (w/v) D-xylose; Ninck et al., unpublished). The *alba* transcript comprises a 14-nucleotide long 5’-UTR (5’ GAU**AGGUG**GUUUAA 3’) (Cohen et al., 2016; Busche et al., unpublished), including a 5-nucleotide long SD sequence motif (bold). To evaluate the impact of this 5’-UTR on protein production, we selected the esterase Saci_1116 as reporter protein. The enzyme was previously characterized (Sobek & Görisch, 1988, 1989), and was used as a model protein in the optimization of the *S. acidocaldarius* expression system before (van der Kolk et al., 2020). We employed existing expression plasmids containing the P_ara_ promoter and encoding *saci_1116*, with either an N- or C-terminal Twin-Strep tag (NtSS or CtSS, respectively; Tab. 1) as templates for *alba* 5’-UTR insertion and as reference (Fig. S1). The *alba* 5’-UTR was inserted upstream of the reporter gene start codon (“ATG”; Fig. S1), resulting in the generation of *S. acidocaldarius* expression plasmids termed pBSaraFX-***alba*UTR**-*saci_1116*-NtSS or -CtSS, respectively (Fig. S1, S2; Tab. 1). Reporter protein production and purification from the soluble protein fraction were evaluated using SDS PAGE (Fig. 2).

**Figure 2:**
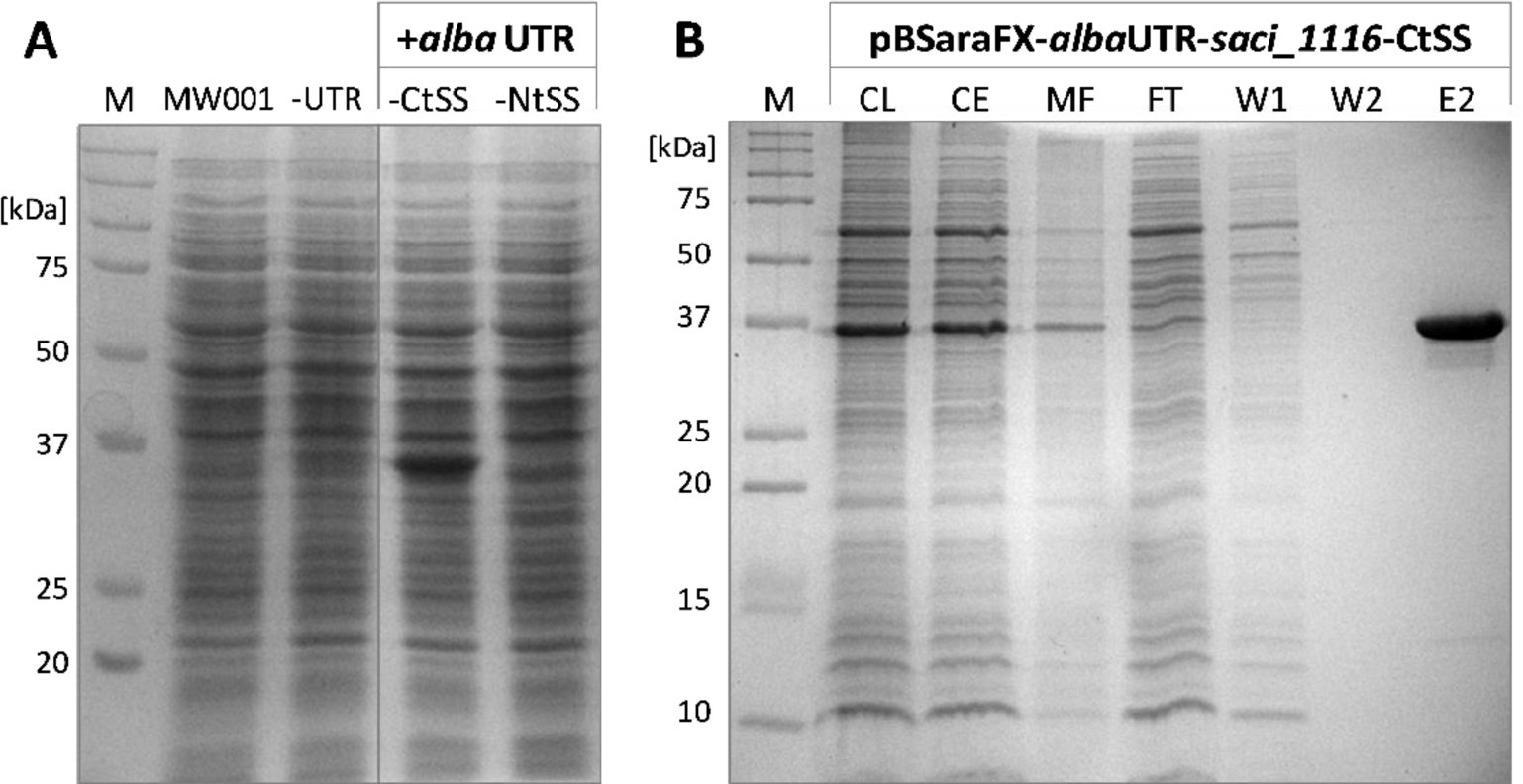
Effect of *alba* 5’-UTR on esterase Saci_1116 production in *S. acidocaldarius* MW001. The expression without and with *alba* 5’-UTR, and with C-terminal (Ct) or N-terminal (Nt) Twin-Strep tag (SS) in Brock medium (pH 3.0) supplemented with 0.1% (w/v) NZA and 0.3 (w/v) D-xylose at 140 rpm and 76°C is shown. *S. acidocaldarius* MW001 without plasmid serves as reference (MW001). -UTR: pSVAaraFX-*saci_1116*-CtSS; +*alba* UTR: pBSaraFX-*alba*UTR-*saci_1116*-CtSS/NtSS. **A)** Coomassie-stained SDS-PAGE gel of cell lysates. For all cell samples, an OD_600nm_ value of 10 was adjusted and 20 µL of each sample were applied to SDS-PAGE. **B)** SDS-PAGE image of Saci_1116-CtSS purification from crude extract of *S. acidocaldarius* MW001 pBSaraFX-*alba*UTR-*saci_1116*-CtSS by affinity chromatography (M: marker, CL: cell lysate; CE: crude extract (soluble fraction); MF: membrane fraction; FT: flow-through; W: wash fraction; E: elution, 3 µg protein). Theoretical molecular weights: Saci_1116-NtSS: 37.1 kDa, Saci_1116-CtSS: 36.8 kDa.

Insertion of the *alba* 5’-UTR into the *saci_1116*-CtSS expression plasmid markedly enhanced reporter protein yield, as evidenced by SDS-PAGE analysis (Fig. 2A). The utilization of an N-terminal tag yielded lower Saci_1116 production in contrast to a C-terminal tag, in line with previous observations (van der Kolk et al., 2020). Affinity chromatography of the soluble protein fraction (crude extract) from the expression culture (*S. acidocaldarius* MW001 pBSaraFX-*alba*UTR-*saci_1116*-CtSS) efficiently purified Saci_1116-CtSS using the C-terminal Twin-Strep tag (Fig. 2B). Only minor amounts of reporter protein were detected in the membrane fraction. Bradford protein quantification revealed a four-fold increase in reporter protein yield from crude extracts using the *alba* 5’-UTR modified expression plasmid compared to the vector lacking the 5’-UTR (Fig. S3). The purified esterase Saci_1116-CtSS exhibited a specific activity of 124.5 U/mg with *p*NP-acetate as substrate at 42°C. In conclusion, incorporation of the *alba* 5’-UTR into the *saci_1116*-CtSS expression plasmid significantly enhanced the production of soluble, active esterase reporter protein.

### Effect of 5’-UTR insertion on promoter activity

To assess the impact of *alba* 5’-UTR on the functionality and selective induction of the pentose-inducible promoter (*saci_2122* promoter P_ara_), we investigated its activity under the influence of different sugars (Fig. S4). Our findings demonstrate that the functionality of the pentose-inducible promoter remained unaffected by the insertion of the *alba* 5’-UTR into the expression construct (Fig. S4). In agreement with previous studies, gene expression was induced by the addition of pentoses, while minimal basal expression was observed with sucrose, the disaccharide comprising D-glucose and D-fructose (Fig. S4; van der Kolk et al., 2020). Thus, *alba* 5’-UTR insertion resulted in a profound increase in reporter protein levels, while maintaining P_ara_’s inducibility and pentose specificity.

### Influence of −2 and −1 nucleotide identities of *alba* 5’-UTR on reporter protein production

Expression vectors should facilitate the seamless integration of genes of interest (GOIs) through appropriate restriction sites and cloning strategies. In pSVAara expression vectors, a *Nco*I restriction site (5’ <u>C’C</u>ATGG 3’) serves as the upstream cloning site in restriction- and ligation-dependent GOI integration. For vectors featuring N-terminal protein tags, this *Nco*I site is positioned downstream of the tag-encoding nucleotide sequence. Thus, the integration of a 5’-UTR upstream of the N-terminal tag-coding sequence does not affect the *Nco*I cloning site (Fig. S1). However, in vectors with C-terminal protein tags, the direct insertion of the “original” *alba* 5’-UTR sequence (5’ GATAGGTGGTTT<u>AA</u> 3’) upstream of the translation start codon (“ATG”) would disrupt the *Nco*I restriction site used for GOI integration, which simultaneously encodes the translation start codon in C-terminal tag vectors (*Nco*I site: 5’ <u>C’C</u>**ATG**G 3’, start codon in bold) (Fig. S1).

Therefore, the two nucleotides located at the 3’-end of the 5’-UTR, immediately upstream of the start codon, hereafter referred to as −2 and −1 nucleotides, were modified to preserve the *Nco*I restriction site in C-terminal protein tag vectors. Specifically, the identities of the −2 and −1 nucleotides were changed from “AA” to “CC” while maintaining the overall length of the *alba* 5’-UTR. We assessed the impact of these nucleotide changes (i.e. from “AA” to “AC” or “CC”) on reporter protein production through enzyme activity assays, SDS-PAGE, and immunodetection (Fig. 3).

**Figure 3:**
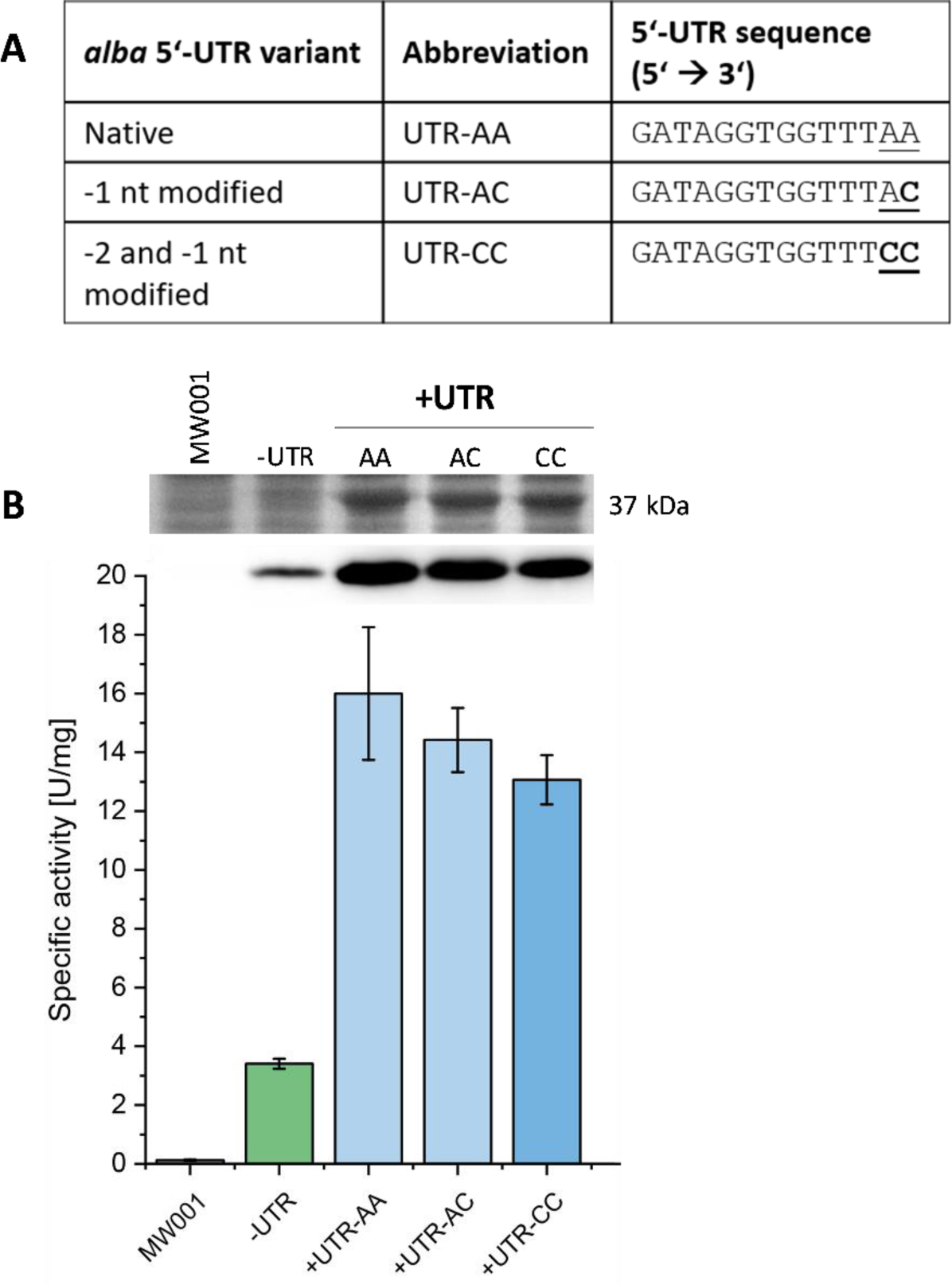
Influence of −2 and −1 nucleotide identities in *alba* 5’-UTR on the production of C-terminal tagged esterase. Expression studies were performed using different *saci_1116*-CtSS expression constructs, either without 5’-UTR (-UTR) or with the *alba* 5’-UTR with different nucleotide sequences at the −2 and −1 positions, i.e. AA, AC, or CC (**A**). Expression cultures were harvested at OD_600nm_ values of 0.6 (log growth phase). *S. acidocaldarius* MW001 without any plasmid served as reference (MW001). **B**) Esterase activity in cell lysate samples of *S. acidocaldarius* MW001 expression cultures as determined in a continuous assay using *p*NP-acetate as substrate, following *p*NP formation at 410 nm and 42°C. Error bars indicate the errors of three biological replicates (*n*=3). Excerpts from respective SDS-PAGE analysis of cell lysate samples (upper lane) and Saci_1116-CtSS immunodetection (lower lane) are shown (for complete images see Fig. S5).

The specific esterase activities determined in cell lysates of *S. acidocaldarius* expression cultures exhibited similarity across all tested −2 and −1 nucleotide identities (“AA”, “AC” and “CC”) (Fig. 3A), as supported by standard deviations and Student’s t-test results (Tab. S2, Tab. S3). SDS-PAGE analysis and esterase immunodetection confirmed the results of activity measurements (Fig. 3B; Fig. S5). In comparison to expression conditions lacking a 5’-UTR, the insertion of the *alba* 5’-UTR variants resulted in an average 4.5-fold increase in esterase activity, with the reporter enzyme constituting approx. 11% of cell protein (Tab. S2).

### Screening of different UTR candidates from *S. acidocaldarius* for efficient reporter protein production

In the next step, we sought to ascertain whether the efficacy of the expression system could be augmented by utilizing alternative 5’-UTR sequences. Leveraging literature and available proteomic data, we identified proteins of notable abundance in *S. acidocaldarius* and extracted their respective 5’-UTR sequences from RNA sequencing data (Weizmann Exploration, https://exploration.weizmann.ac.il/TCOL/; Cohen et al., 2016). Four 5’-UTR sequences, derived from genes *saci_0330* (encoding a hypothetical protein), *saci_1401* (thermosome α subunit), *saci_2355* (surface-layer protein A), and *saci_2354* (S-layer protein B), were chosen for comparative analysis alongside the *alba* 5’-UTR (Fig. 4A).

**Figure 4:**
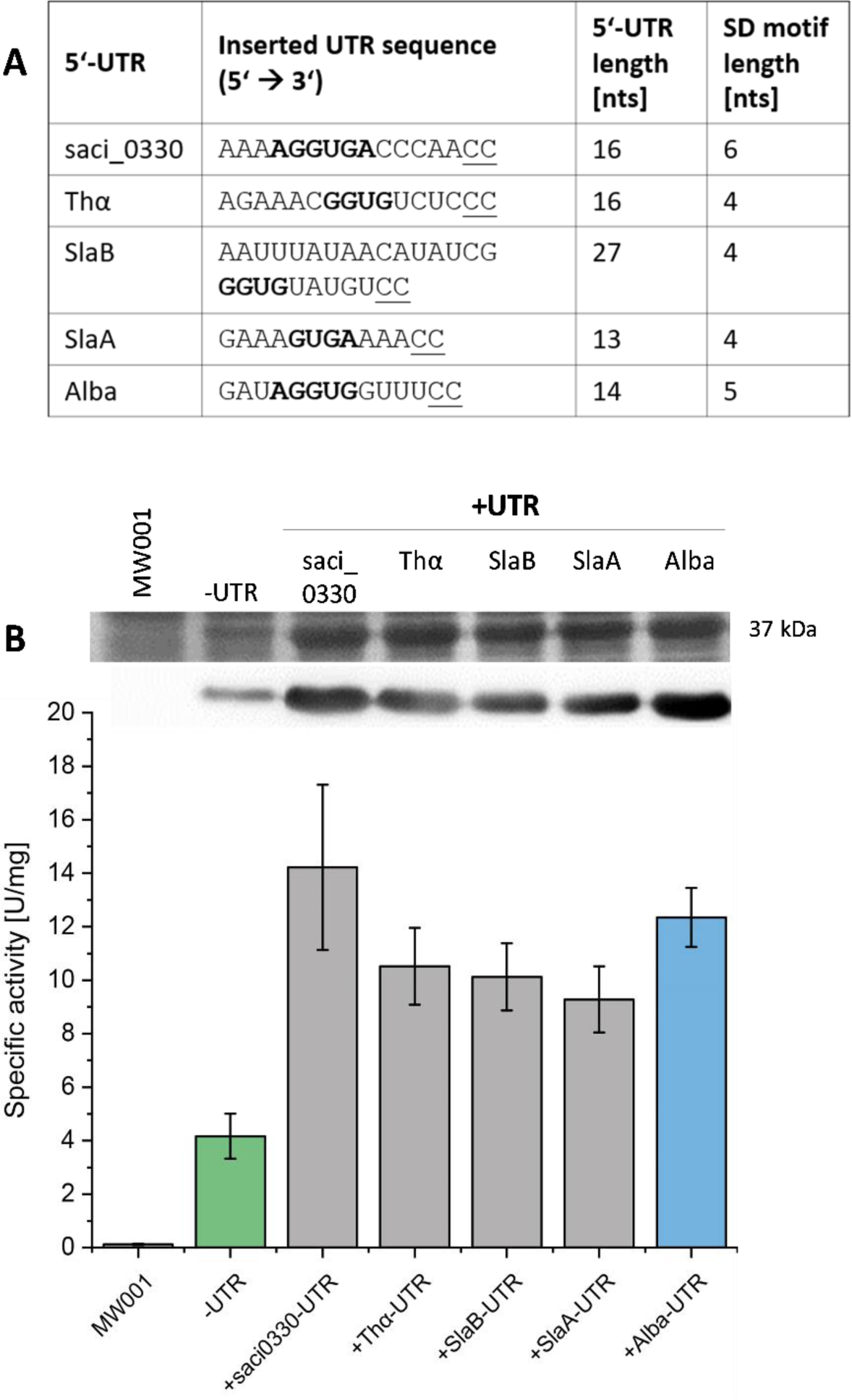
Screening of five different 5’-UTR sequences for the production of Saci_1116-CtSS. **A)** Overview of tested 5’-UTR sequences. Gene IDs, 5’-UTR sequences with their respective SD motifs (bold), UTR lengths, and SD motif lengths are given. **B)** Saci_1116-CtSS production was followed using constructs without (-UTR) or with a 5’-UTR (+UTR). 5’-UTR sequences of genes *saci_0330*, *saci_1401* (Thα), *saci_2354* (SlaB), *saci_2355* (SlaA) and *saci_1322* (Alba) were inserted into the *saci_1116*-CtSS expression plasmid directly upstream of the translation start codon (Tab. 1). Cultivation and protein quantification were performed as described in figure legend 2. Expression cultures were harvested at OD_600nm_ values of 0.8−1.2 (end of log growth phase). Specific esterase activity was determined for cell lysates using the continuous *p*NPA assay (*n*=3). Excerpts from respective SDS-PAGE analysis of cell lysate samples (upper lane) and Saci_1116-CtSS immunodetection (lower lane) are shown (for complete images see Fig. S6).

The 5’-UTR sequence of *saci_0330* corresponds to the transcript leader sequence of the operon *saci_0330-0333*, with *saci_0331* annotated to encode pyridine nucleotide-disulphide oxidoreductase, while the others are annotated as conserved proteins of unknown functions (Cohen et al., 2016). The 5’-UTR sequences of *saci_2355* and *saci_2354,* encoding S-layer proteins A and B, respectively, are encoded in a single operon (*slaAB*) on the complementary DNA strand (Veith et al., 2009). However, in addition to the promoter upstream of SlaA, available transcriptome data revealed a second promoter and a weak 5’-UTR signal upstream of *slaB*, consistent with RNA sequencing data (Cohen et al., 2016). The 5’-UTR of *saci_1401*, the gene encoding the thermosome α subunit (*thα*), a major chaperone complex in Sulfolobales (Chaston et al., 2016; Baes et al., 2020), and the *alba* 5’-UTR originate from single gene transcripts (Cohen et al., 2016). 5’-UTRs of *saci_0330* and *alba* contained 6- and 5-nucleotide long SD motifs, respectively, while the other three sequences (*thα*, *slaB, slaA*) included SD motifs of 4 nts in length (Fig. 4A).

As before, the *Nco*I restriction and cloning site (5’ <u>C’C</u>**ATG**G 3’) was maintained, i.e., the nucleotide identities at positions −2 and −1 (relative to the translation start codon) were changed to “CC” for all 5’-UTR sequences tested (Fig. 4A, underlined), while preserving the overall length of each 5’-UTR (Tab. S4). Screening was conducted as previously described using esterase Saci_1116-CtSS as reporter protein, and its production was evaluated by specific esterase activity, SDS-PAGE, and immunodetection (Fig. 4).

The 5’-UTR screening revealed significantly increased reporter protein activity in the presence of all tested 5’-UTR sequences compared to the absence of a 5’-UTR (Fig. 4; Tab. S5, S6). Confirmation of these activity-based results was obtained through SDS-PAGE analysis and esterase immunodetection (Fig. 4B; Fig. S6). Integration of different 5’-UTR sequences increased the amount of reporter protein to approx. 7.5−11% of the total protein compared to 3% without an inserted 5’-UTR (Tab. S5). On average, *saci_0330* and *alba* 5’-UTRs demonstrated the highest production yield of active esterase (Fig. 4B). However, the *alba* 5’-UTR exhibited the lowest standard deviations among biological replicates and the smallest p-value when comparing esterase activities with and without a 5’-UTR (Tab. S6).

To ensure generalizability regarding the impact of the screened 5’-UTRs on different proteins, we also investigated their impact on another reporter protein, β-galactosidase LacS from *S. solfataricus* (Fig. S7) (Pisani et al., 1990). LacS was commonly employed as a reporter in various studies (e.g. Jonuscheit et al., 2003; Wagner et al., 2012), including promoter investigations (Berkner et al., 2007; 2010; Peng et al., 2012; van der Kolk et al., 2020) or analyses of oligosaccharide hydrolysis (e.g. Curci et al., 2021). In contrast to esterase Saci_1116, LacS exhibits substantial expression levels even without 5’-UTR. Nevertheless, the addition of 5’-UTRs resulted in further enhancement of LacS production. Specifically, the *alba* 5’-UTR led to a 1.6-fold increase in LacS production (Fig. S7). Remarkably, the effect of 5’-UTRs on target protein production exhibited a general, consistent pattern, demonstrating the reproducibility of the observed effects. However, the factor of protein production enhancement by the integration of 5’-UTRs is target protein-dependent, as shown for LacS.

### Application of the *alba* 5’-UTR optimized expression vector for the production of glycosyltransferases

Our previous experiments illustrated the effectiveness of 5’-UTR-optimized protein production for two different target proteins. To further investigate the transferability of the newly modified expression vectors, we considered their impact on challenging proteins as particularly relevant. In several studies, the production of glycosyltransferases (GTs) has been reported as demanding (e.g. Breton et al., 2006; Cobucci-Ponzano et al., 2011; Mestrom et al., 2019; MoseRossi et al., 2010). Accordingly, also GTs originating from *S. acidocaldarius* have proven difficult to express in both *E. coli* and *S. acidocaldarius* when using vectors lacking a 5’-UTR. For instance, the GT-encoding gene s*aci_1911* exhibited weak expression in *E. coli* (pET15b, N-terminal His_6_-tag) and appeared to form insoluble aggregates (data not shown). Thus, we utilized the new *alba* 5’-UTR modified expression vectors, designated as **pBSaraFX-*alba*UTR-NtSS and -CtSS** (Fig. S8, S9), for GT synthesis in *S. acidocaldarius*, assessing the production yield in comparison to constructs lacking the *alba* 5’-UTR (Fig. 5). Saci_1911-CtSS was purified from crude extracts of expression cultures by affinity chromatography, and proteins were visualized by SDS-PAGE (Fig. 5).

**Figure 5:**
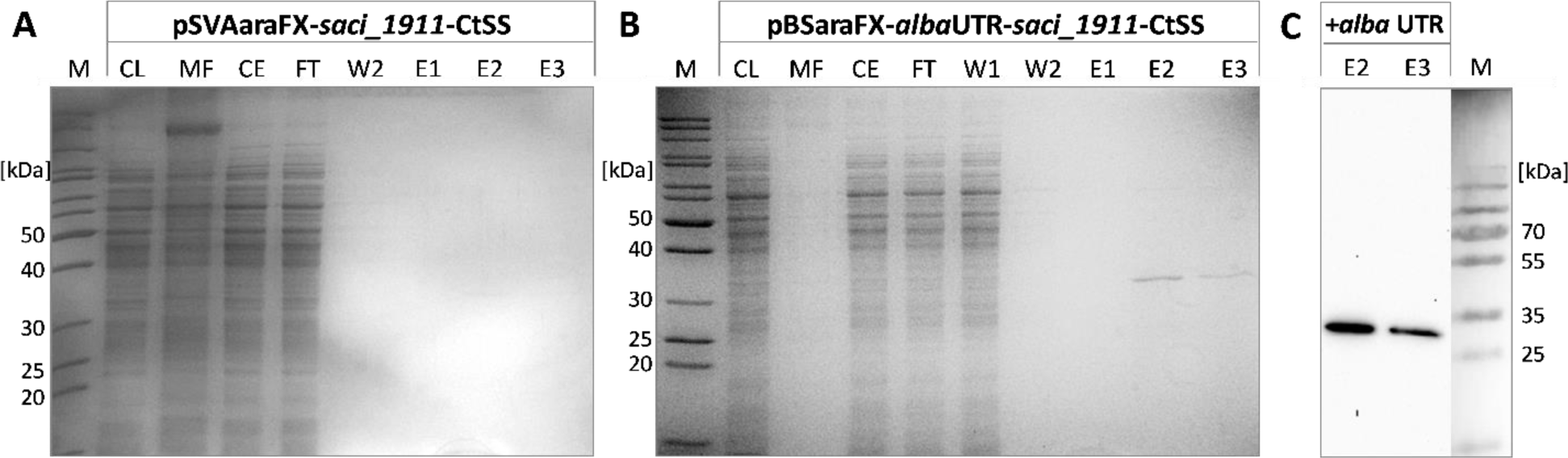
Effect of the *alba* 5’-UTR on the yield of homogenously produced and purified glycosyltransferase Saci_1911-CtSS. SDS-PAGE gels of Saci_1911-CtSS affinity chromatography fractions after expression in *S. acidocaldarius* MW001 using pSVAaraFX-*saci_1911*-CtSS (no 5’-UTR) (**A**) and pBSaraFX-*alba*UTR-*saci_1911*-CtSS (+*alba* 5’-UTR (**B**). **C**) Immunodetection of Twin-Strep-tagged Saci_1911-CtSS in affinity chromatography elution fractions (+*alba* UTR) using Strep-Tactin-HRP conjugate. Expression cultures (150 mL each) were cultivated as described in the legend of figure S3 and harvested at OD_600nm_ values of 0.8. Protein was purified from crude extracts (of 0.3 g cell wet weight each) using the Strep-Tactin®XT Superflow® Kit following the manufacturer’s protocol (200 µL column volume each). Theoretical MW of Saci_1911-CtSS: 33.8 kDa (M: marker; CL: cell lysate; MF: membrane fraction (pellet after cell lysis); CE: crude extract; FT: flow-through fraction of affinity chromatography; W: wash fraction; E: elution fraction, 20 µL each (0.5 µg Saci_1911-CtSS in E2 of pBSaraFX-*alba*UTR-*saci_1911*-CtSS (B)).

**Figure 6:**
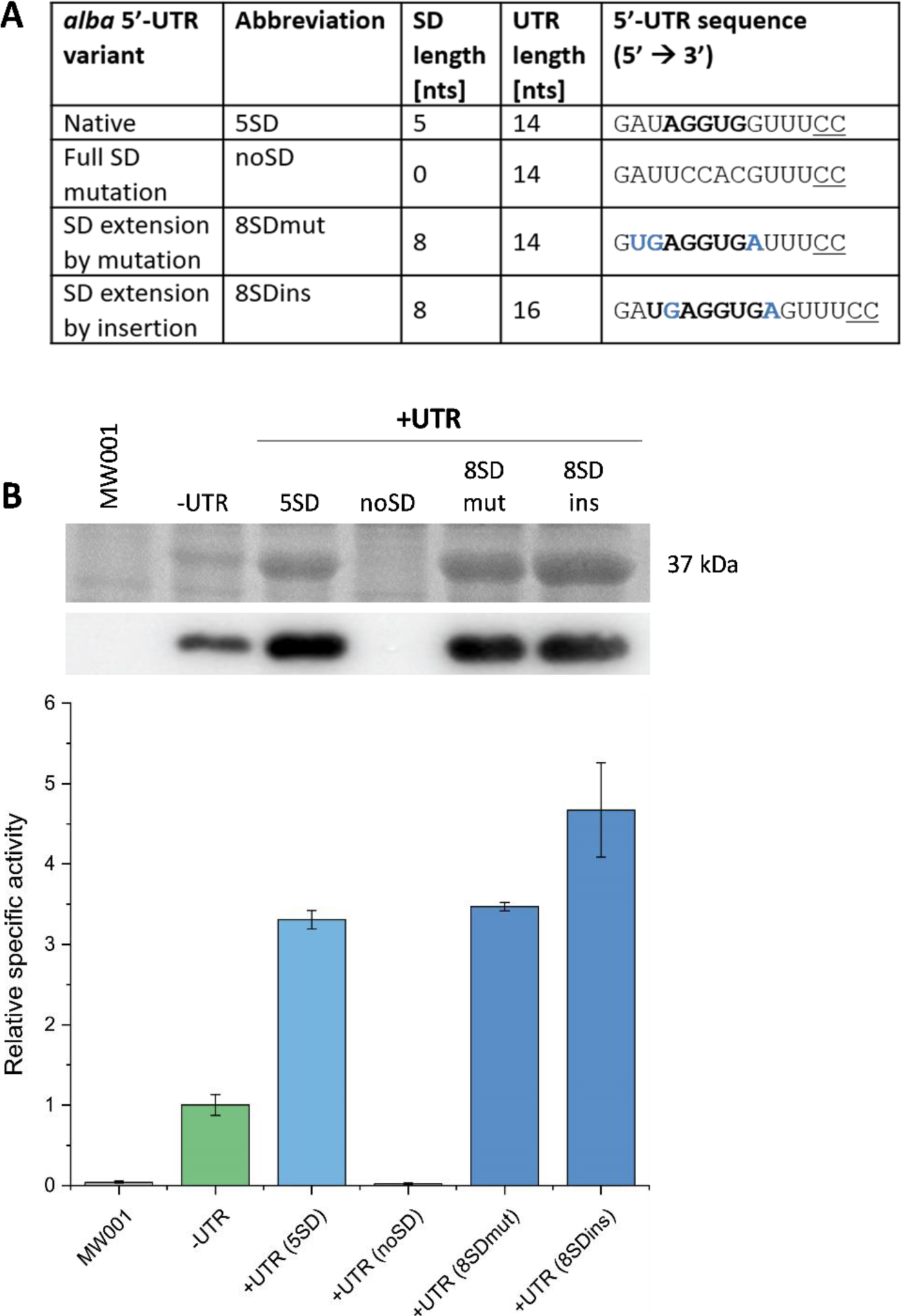
Effect of SD motif sequence alterations within the *alba* 5’-UTR on esterase production. **A**) Structural properties of tested *alba* 5’-UTR-SD variants. SD lengths, UTR lengths, and 5’-UTR nucleotide sequences are given. **B**) Esterase activity in cell lysate samples of *S. acidocaldarius* MW001 expression cultures as determined in a continuous assay using *p*NP-acetate (*p*NP formation detected at 410 nm and 42°C). Error bars indicate the errors of three biological replicates (*n*=3). Excerpts from respective SDS-PAGE analysis of cell lysate samples (upper lane) and Saci_1116-CtSS immunodetection (lower excerpt) are shown (for complete images see Fig. S11). Expression cultures were harvested at OD_600nm_ values of 0.6-0.8 (log growth phase).

When employing the *alba* 5’-UTR modified expression vector and a C-terminal Twin-Strep tag, Saci_1911 was successfully produced in the soluble fraction of *S. acidocaldarius*. Notably, Saci_1911 production was higher with a C-terminal compared to an N-terminal tag (data not shown). While no purified protein was obtained by SDS-PAGE analysis in elution fractions of expression cultures using the vector without 5’-UTR, the GT was successfully synthesized and purified from cells containing the *alba* 5’-UTR expression plasmid (Fig. 5). Thus, from this example, it is evident that the application of the 5’-UTR modified vector can increase yields of challenging proteins, shifting from “no protein” (6 µg from 1 g cell wet weight) to “some” protein (35 µg from 1 g cell wet weight). A similar effect was observed for GT Saci_1915 (Fig. S10). Homologous production of GT Saci_1915, both with an N- or C-terminal Twin-Strep tag and without and with the *alba* 5’-UTR, showed enhanced protein production for both tagged variants using the *alba* 5’-UTR, although an N-terminal tag location was preferred in the case of this GT (Fig. S10).

In summary, the 5’-UTR optimized expression vector successfully enabled or increased the production of different proteins, including glycosyltransferases that were “hard to produce” in the bacterial expression host *E. coli*.

### Impact of Shine-Dalgarno (SD) motif on protein production

As previously noted, the native *alba* 5’-UTR contains a 5 nt long SD motif. This prompted us to investigate the relevance of this motif for efficient protein production. Considering the aSD sequence found in *S. acidocaldarius* 16S rRNA, a SD motif with the sequence “UGAGGUGA” would perfectly complement the aSD over a length of 8 nucleotides (Ma et al., 2002; Schmitt et al., 2020; Wen et al., 2021). To explore the potential enhancement of protein production by extending the SD motif, we constructed *alba* 5’-UTR variants containing 8-nucleotide long SD motifs. The first variant, designated “8SDmut” was generated by site-directed mutagenesis to alter nucleotides neighboring the native 5 nt long SD motif while maintaining the overall UTR length (Fig. 6A). The second variant, named “8SDins”, was generated by the insertion of two nucleotides to create an 8-nucleotide long SD motif, thereby increasing the total length of the 5’-UTR. Additionally, the native 5 nt long SD motif was completely mutated to generate a 5’-UTR lacking the SD motif (“noSD” variant) (Fig. 6A). Throughout these modifications, the overall GC content of the 5’-UTR was kept constant at 50%. Esterase Saci_1116-CtSS was utilized as reporter protein and its production was assessed by enzymatic activity, SDS-PAGE, and immunodetection (Fig. 6).

Strikingly, the absence of the SD motif within the *alba* 5’-UTR resulted in a complete cessation of protein production, as confirmed by enzymatic measurements and protein visualization (Fig. 6). One of the 8-nucleotide long SD variants, “8SDins”, facilitated a further increase in protein production compared to the native *alba* 5’-UTR with a 5-nucleotide long SD motif. In contrast, the variant “8SDmut”, generated through nucleotide mutagenesis, displayed protein production levels comparable to those of the native *alba* 5’-UTR (Fig. 6). These results suggest that the sole extension of the SD motif does not account for the observed increase in protein production, indicating the presence of complex effects at the sequence or potentially transcript level. Importantly, our findings underscore the crucial role of the SD motif within the *alba* 5’-UTR in allowing protein expression.

## 4 Discussion

In this study, we examined the impact of 5’-UTR sequence insertions into *S. acidocaldarius* P_ara_ expression plasmids on the production of different (reporter) proteins. Our findings elucidate that the integration of SD motif-containing 5’-UTRs from proteins with high cellular abundance significantly enhanced plasmid-based protein synthesis for all tested target proteins.

The utilization of 5’-UTRs and RBS, i.e. preserved Shine-Dalgarno sequences, is prevalent in bacterial expression vectors, such as pBAD, pET, pMAL, pQE, and pGEX plasmids for heterologous expression in *E. coli,* or pHT and pBSMul1 expression plasmids in *Bacillus subtilis* (Brockmeier et al., 2006; Yang et al., 2021). Consequently, the influence of bacterial 5’-UTR sequence elements, i.e. SD motifs or spacer sequences, has been extensively investigated in several bacterial organisms over the last decades (e.g. Wu & Janssen, 1997; Sakai et al., 2001; Khosa et al., 2018; Volkenborn et al., 2020). In contrast, 5’-UTRs have been less considered and studied as a relevant feature for gene overexpression in Archaea. However, this disparity is likely related to the fact that, unlike bacteria, many common archaeal model organisms, including Haloarchaea, Thermoproteales, and Sulfolobales, primarily generate leaderless gene transcripts (e.g. 72% for *H. volcanii* (Schmitt et al., 2020), or 69% for *S. solfataricus* (Wurtzel et al., 2010); Slupska et al., 2001; Brenneis et al., 2007, Kramer et al., 2014). Consequently, 5’-UTRs were considered less relevant for protein synthesis and as a feature of archaeal expression vectors. However, some existing archaeal expression vectors do generate leadered mRNA transcripts for gene overexpression, facilitated by the presence of an extended promoter sequence that generates a 5’-UTR. For instance, the sequence of the heat-inducible tf55 promoter in *S. solfataricus* expression vectors generates an 18 nt long 5’-UTR harbouring a putative 4 nt long SD motif (Jonuscheit et al., 2003). The existing sugar-inducible *S. acidocaldarius* expression vectors featuring P_ara_ from *saci_2122* and P_mal_ (maltose) from *saci_1165* generate leaderless transcripts, as determined by available RNA sequencing data of corresponding gene transcripts from the genome (Cohen et al., 2016). However, in the case of P_xyl_ (promoter from *saci_1938*) (van der Kolk et al., 2020), a leadered transcript devoid of an SD motif is generated (34 nts, Cohen et al., 2016).

Here, we introduced SD motif-containing 5’-UTR sequences from genes encoding highly abundant proteins into the P_ara_ *S. acidocaldarius* expression vector. All five tested 5’-UTR insertions demonstrated enhanced production of esterase Saci_1116, showing a two- to four-fold increase. Notably, the use of *alba* (*saci_1322*) 5’-UTR, encoding the most abundant protein in *S. acidocaldarius* according to full proteome data (Ninck et al., unpublished), resulted in a four-fold increase in the yield of soluble, active esterase, along with an additional enhancement in LacS production. Accordingly, in *Methanococcus maripaludis* the replacement of a wild-type 5’-UTR by 5’-UTRs from highly expressed genes (e.g. *slp* (S-layer protein) and *hmmA* (histone A)) led to increased protein production (Akinyemi et al., 2021). Thus, both in *M. maripaludis* and in *S. acidocaldarius* the utilization of 5’-UTRs originating from highly expressed genes resulted in higher plasmid-based target protein production. In contrast, tested native 5’-UTRs from *H. volcanii* (*hlr* (hoxA-like transcriptional regulator) and *hp* (conserved hypothetical protein)) without an SD motif reduced reporter protein amounts by approx. two-fold when compared to leaderless transcripts (Brenneis & Soppa, 2009). However, two out of four artificial 5’-UTR sequences (20 nts, random sequence, no SD) increased reporter activity by factors of two and five, respectively, in *H. volcanii* (Brenneis et al., 2007; Hering et al., 2009). Moreover, 5’-UTR sequences from genes involved in gas vesicle formation in *Halobacterium salinarum* and a short 5’-terminal nucleotide sequence (4 nts) from heat shock protein 70 (*hsp70*) of *Natrinema* sp. J7 decreased reporter production in *H. volcanii* (Born & Pfeifer, 2019; Chen et al., 2015).

Notably, modifications to the nucleotide identities at positions “−1” and “−2” within the short *hsp70* 5’-UTR revealed varied effects on protein production in *H. volcanii*. For instance, altering the −2 nucleotide from its native “C” to “G” completely abolished protein production, while mutations from “C” to “A” or “U” increased it (Chen et al., 2015). Similarly, in *S. islandicus*, mutations of the two nucleotides upstream of the translation start codon (from “AU” to “CC”) within the 6 nt long 5’-UTR of P_araS_ (Peng et al., 2009) reduced protein levels by approx. 30% (Ao et al., 2013). In our study, we also made changes to the two nucleotides immediately upstream of the translation start codon within the *alba* 5’-UTR to retain the *Nco*I cloning site. However, we did not observe any negative effect of this modification on esterase production. Thus, the impact of the nucleotide identities directly upstream of the translation start codon is likely unique to the overall 5’-UTR sequence and its effects on different regulatory levels, such as transcript stability or translation initiation, rendering it complex to predict. Further investigations, including the determination of transcript levels and stabilities, will help to shed more light into these aspects in the future.

Occasionally, the relevance of the SD (RBS) sequence on protein production has been investigated for some 5’-UTRs in Archaea. Concerning Sulfolobales, previous studies have investigated SD mutations or insertions in *S. solfataricus* and *S. islandicus*, respectively. For *S. solfataricus*, the substitution of two nucleotides of the SD motifs of ORF104 or ORF143 of a bicistronic mRNA (8 and 7 nt long SD motifs, respectively) entirely abolished their translation in a cell-free *in vitro* translational system (Condó et al., 1999). Consistent with this finding, we observed a cessation of protein production upon complete mutagenesis of the SD motif within the *alba* 5’-UTR in *S. acidocaldarius*. The insertion of 14 nts, including an 8 nt long SD motif, into an existing 6 nt long 5’-UTR sequence in P_araS_ expression vectors led to a three-fold increase in reporter activity (β-galactosidase) in *S. islandicus* (Peng et al., 2012; Peng et al., 2017). Similarly, we found that the insertion of two additional nucleotides to the *alba* 5’-UTR, thereby extending the length of the SD motif and the overall UTR length, increased the yield of esterase. In *H. volcanii*, mutations to the 7-nucleotide SD motif within the *sod2* 5’-UTR had no impact on reporter protein production, but an extension to 8 nts reduced translational efficiency to approximately 10% (Kramer et al., 2014). For the *gvpH* upstream region from *H. salinarum* mutations to its 7 nt long SD motif resulted in varied reductions in reporter protein production, with a full SD mutation retaining 20% of the original activity (Sartorius-Neef & Pfeifer, 2004). In *T. kodakarensis*, mutations to the second nucleotide of a 6-nucleotide SD motif significantly decreased protein production (Santangelo et al., 2008). Concludingly, the relevance of SD motifs within archaeal 5’-UTRs remains uncertain, emphasizing the need to further investigate their impact on protein expression at the different regulatory levels.

Altogether, reports on the influence of 5’-UTRs in archaea suggest that, compared to leaderless transcripts, 5’-UTRs do not consistently enhance protein production. Multiple factors may contribute to the differential effects of 5’-UTR sequences on protein production efficiency, including complex regulatory mechanisms at the posttranscriptional level (e.g. secondary structures, transcript stability), as well as alterations to the ribosome docking site for translation initiation. This complexity likely applies to *S. acidocaldarius* as well; however, our selection of 5’-UTR sequences from genes with high protein abundance probably precluded the appearance of 5’-UTRs that might reduce protein production.

### Conclusion

For many archaeal genes, heterologous expression in mesophilic bacterial strains, such as *E. coli*, is effective, with established straightforward purification protocols, including heat precipitation, for thermophilic proteins. However, there are numerous examples where attempts to express archaeal proteins in mesophilic bacterial hosts fail. Thus, archaeal expression systems are needed for synthesizing demanding archaeal proteins, often with biotechnological potential, that cannot be efficiently produced in bacterial hosts. Our study showcases the optimization of plasmid-based protein production through the incorporation of SD motif-containing 5’-UTR sequences in the archaeal expression host *S. acidocaldarius*. The integration of all tested 5’-terminal sequences into P_ara_ expression vectors led to a significant enhancement in protein yield, exemplified by the substantial increase in the production of esterase Saci_1116. Notably, among the five different 5’-terminal sequences examined, the 5’-UTR of the Alba-encoding gene *saci_1322* was found to be highly efficient and reliable. The *alba* 5’-UTR containing expression vector proved to be effective for the synthesis of challenging target proteins, as shown for archaeal, thermophilic glycosyltransferases. Overall, the application of 5’-UTR-optimized expression vectors holds promise for facilitating the synthesis of other challenging thermostable proteins in the future.

## 6 Author Contributions

LK and AW conducted experiments, performed data analysis, and contributed to data visualization. TB and JK, part of the HotAcidFACTORY consortium, aided in determining 5’-UTRs through transcriptomics analysis. CB contributed to data interpretation and manuscript drafting, while CB and BS acquired funding. LK and BS serve as corresponding authors and contributed to experimental design, data interpretation, and manuscript preparation. All authors contributed to the article and approved the submitted version.

## 8 Acknowledgments

We would like to express our sincere gratitude to Malin Kretzschmar and Semira Gertchen for their valuable contributions to this research project.

ChatGPT version 3.5 (OpenAI) was applied for language optimization and to improve the clarity of the scientific content presented.

## 9 Supplementary material

Separate document

