## Supplementary information for "5’-untranslated region sequences enhance plasmid-based protein production in *Sulfolobus acidocaldarius*"

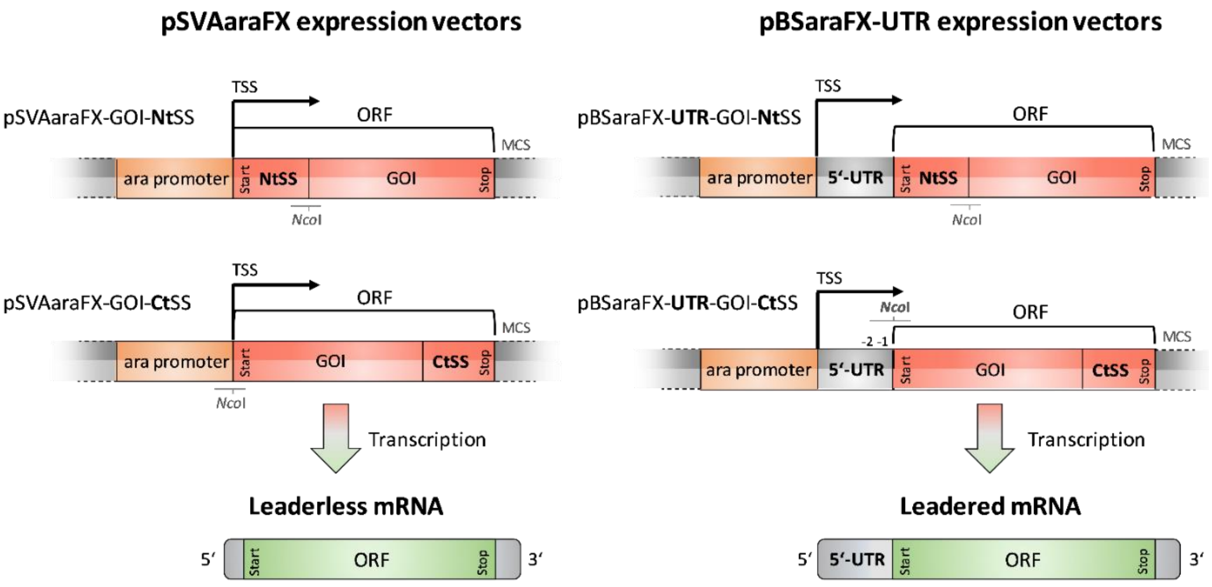


**Figure S1**: Structural elements of pSVAaraFX- and pBSaraFX-UTR expression cassettes and the generation of leaderless or leadered mRNAs, respectively. Upper row: Twin-Strep tag expression vectors with an N-terminal tag location; lower row: C-terminal tag location. The position of the *Nco*I restriction site and of -2 and -1 nucleotides (upstream of the translation start codon) within the 5'-UTR of pBSaraFX-UTR-GOI-CtSS are indicated. ara promoter: *saci_2122* promoter; TSS: transcription start site; ORF: open reading frame; NtSS: N-terminal Twin-Strep tag; GOI: gene of interest; CtSS: C-terminal Twin-Strep tag; Start: translation start codon; Stop: translation stop codon; MCS: multiple cloning site.

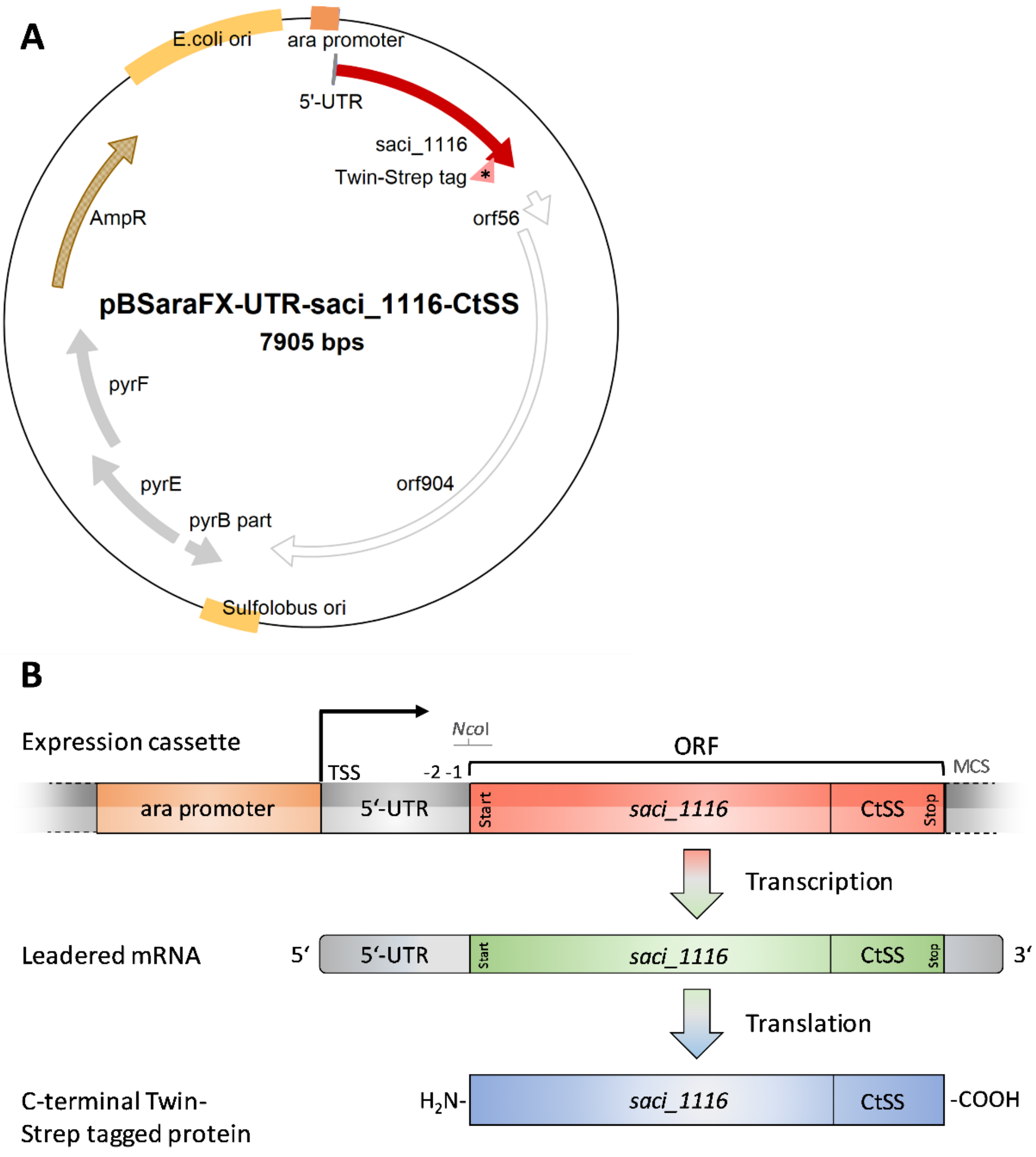


**Figure S2**: **A**) Map of pBSaraFX-*alba*UTR-*saci_1116*-CtSS. The gene of interest, *saci_1116*, was cloned in-frame under control of the pentose-inducible promoter (promoter of *saci_2122*). A 5'-UTR sequence, here the *alba* 5'-UTR, was cloned directly upstream of the translation start codon (ATG). A C-terminal Twin-Strep tag (-CtSS) coding sequence was added to the C-terminal end of the GOI (reporter gene *saci_1116*). The asterisk within the Twin-Strep tag coding element indicates that this sequence location is variable (downstream of the GOI in -CtSS constructs, upstream of the GOI in -NtSS expression constructs) (Fig. S1). Classical shuttle vector elements, such as an origin of replication (ori) for *E. coli* and *S. acidocaldarius*, the ampicillin resistance cassette (AmpR) for selection in *E. coli* as well as *pyrEF* and part of *pyrB* (containing the *pyrEF* promoter sequence) from *Saccharolobus solfataricus* for selection in *S. acidocaldarius* MW001 are indicated. *orf56*, encoding a DNA-binding protein, and *orf904*, encoding a multifunctional replication protein, originate from plasmid pRN1 from *S. islandicus* and regulate plasmid copy number (Lipps, 2004). Clone Manager 7 (Sci Ed Software, USA) was used to generate the figure. **B**) Generation of leadered mRNA and C-terminal (-Ct) Twin-Strep-tagged (SS) protein from the expression cassette present in pBSaraFX-(*alba*)UTR-*saci_1116*-CtSS plasmids. The location of *Nco*I restriction site, as well as -2 and -1 nucleotides within the 5'-UTR sequence are indicated. ara promoter: *saci_2122* promoter; TSS: transcription start site; ORF: open reading frame; CtSS: C-terminal Twin-Strep tag; MCS: multiple cloning site.

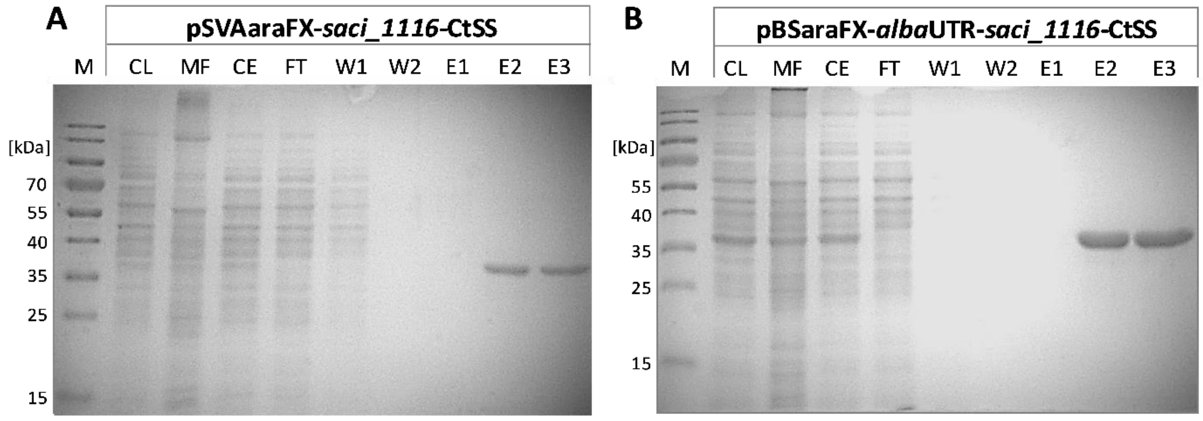


**Figure S3**: Effect of *alba* 5'-UTR on homologous production of the esterase reporter protein Saci_1116-CtSS. Coomassie-stained SDS-PAGE gel of purified Saci_1116-CtSS after expression in *S. acidocaldarius* MW001 using pSVAaraFX-*saci_1116*-CtSS (**no UTR; A**) and pBSaraFX-*alba*UTR-*saci_1116*-CtSS (**+*alba*UTR; B**). Expression cultures were grown under the same growth conditions in Brock medium (200 mL each, pH 3.0) supplemented with 0.1% (w/v) NZA and 0.3 (w/v) D-xylose at 140 rpm and 76°C and harvested simultaneously at $OD_{600nm}$ values of 1.0 and 0.9, respectively. A theoretical turbidity ($OD_{600}$) of 5 was applied for cell lysis by sonication. Protein purification from crude extracts (soluble fraction) was performed using the Strep-Tactin®XT Superflow® Kit according to the manufacturer's protocol. (M: marker (protein ladder); CL: cell lysate (whole cell sample after lysis); CE: crude extract (soluble fraction); MF: membrane fraction (pellet after cell lysis); FT: flow-through fraction of affinity chromatography; W: wash fraction; E: elution fraction, 15 µL each). Theoretical MW of Saci_1116-CtSS: 36.8 kDa.

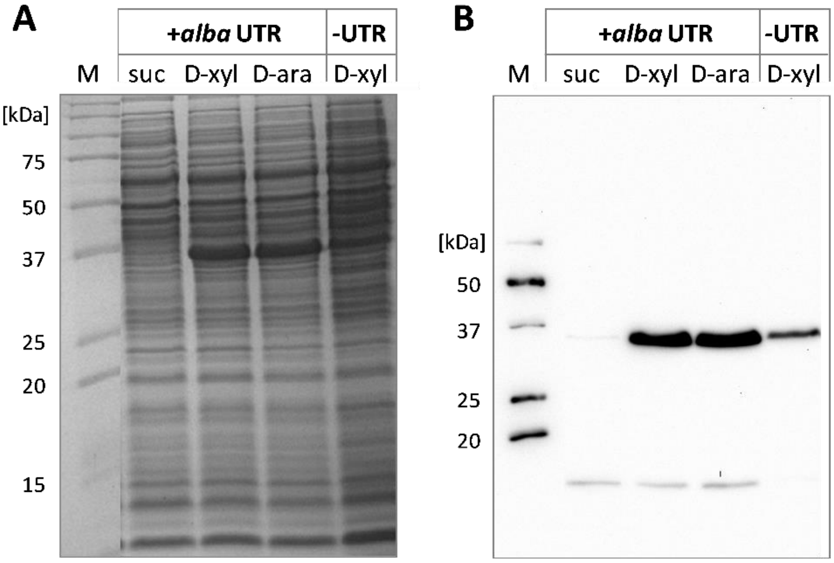


**Figure S4**: Effect of *alba* 5'-UTR insertion on promoter functionality and activity. *S. acidocaldarius* MW001 was transformed with pBSaraFX-*alba*UTR-*saci_1116*-CtSS (+*alba* UTR) or pSVAaraFX-*saci_1116*-CtSS (-UTR). Expression cultures were grown in Brock medium (pH 3.0) with 0.1% (w/v) NZA and different carbon sources (suc: 0.3% (w/v) sucrose; D-xyl: 0.3% (w/v) D-xylose, D-ara: 0.05% (w/v) D-arabinose) at 76°C. A) Coomassie-stained SDS-PAGE gel of cell lysate samples. B) Immunodetection of the Twin-Strep-tagged esterase Saci_1116 using Strep-Tactin-HRP conjugate. Theoretical MW of Saci_1116-CtSS: 36.8 kDa. M: marker (protein ladder).

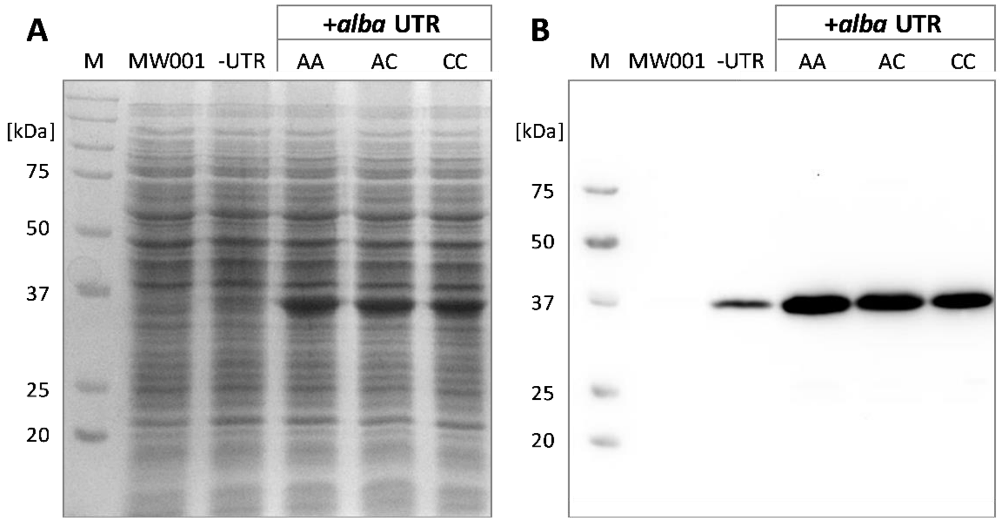


**Figure S5**: Influence of -2 and -1 nucleotide identities in *alba* 5'-UTR on C-terminal tagged esterase reporter protein production. Expression studies were performed using different *saci_1116*-CtSS expression constructs, either without 5'-UTR (-UTR) or with the *alba* (*saci_1322*) 5'-UTR (+*alba* UTR) with different nucleotide sequences at the -2 and -1 positions, i.e. AA, AC or CC. Saci_1116-CtSS production was visualized by SDS-PAGE and Coomassie staining (**A**) and immunodetection (Strep-Tactin HRP conjugate) (**B**). Cultivation and protein quantification were performed as described in the legend of Fig. 3 (main text). M: marker (protein ladder).

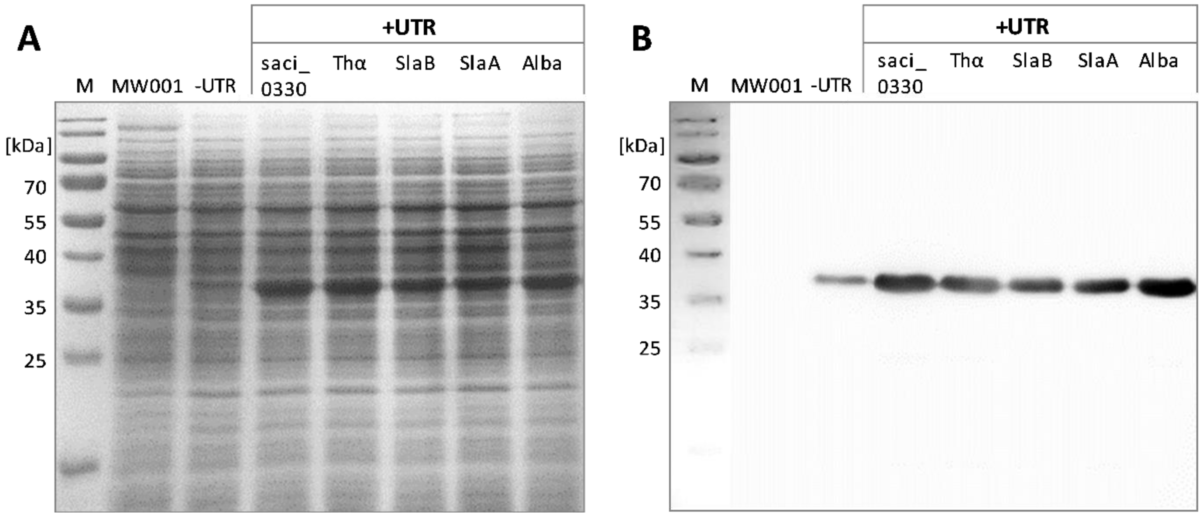


**Figure S6**: Screening of five different 5'-UTR sequences for the production of C-terminal tagged esterase reporter protein (Saci_1116-CtSS). **A**) Visualization of proteins in cell lysates by SDS-PAGE and Coomassie staining. **B**) Immunodetection of C-terminal tagged esterase in cell lysate samples (Strep-Tactin HRP conjugate). Saci_1116-CtSS production was performed using constructs without (-UTR) or with a 5'-UTR (**+UTR**). 5'-UTR sequences of genes *saci_0330*, *saci_1401* (Thα), *saci_2354* (SlaB), *saci_2355* (SlaA) and *saci_1322* (Alba), respectively, were modified in favor of the *Nco*I site (Tab. S4) and inserted into the *saci_1116*-CtSS expression plasmid directly upstream of the GOI start codon. Cultivation and protein quantification were performed as described in the legend of Fig. 4 (main text). M: marker (protein ladder).

**A**

| 5'-UTR | Inserted 5'-UTR sequence (5' → 3') | UTR length [nts] | SD length [nts] | Spacer length [nts] | Spacer GC content [%] | UTR GC content [%] |
|---|---|---|---|---|---|---|
| saci_0330 | AAA**AGGUGA**CCCAACC | 16 | 6 | 8 | 63 | 50 |
| Thα | AGAAAC**GGUG**UCUCCC | 16 | 4 | 6 | 67 | 56 |
| SlaB | AAUUUAUAACAUAUCG**GGUG**UAUGUCC | 27 | 4 | 7 | 43 | 33 |
| SlaA | GAAA**GUGA**AAACC | 13 | 4 | 6 | 33 | 38 |
| Alba | GAU**AGGUG**GUUUCC | 14 | 5 | 6 | 50 | 50 |

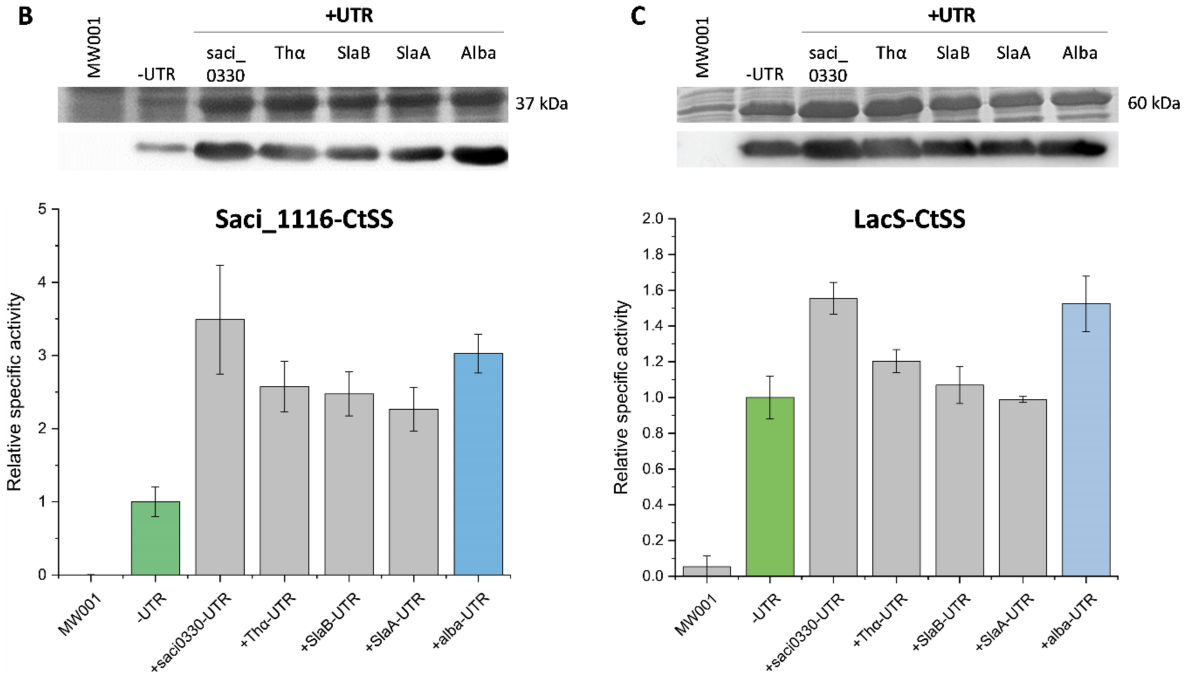


**Figure S7**: **Screening of 5'-UTR sequences for the production of two distinct reporter proteins. A**) Structural properties of tested 5'-UTR sequences. The table presents the structural properties of the tested 5'-UTR sequences, including gene IDs and nucleotide changes at positions -2 and -1 (to "CC", underlined), presence of Shine-Dalgarno motifs (highlighted in bold), UTR lengths, putative SD motif lengths, spacer lengths, and GC contents of spacers and total UTR sequences. **B**) Esterase Saci_1116-CtSS production with different 5'-UTRs. Specific activity of Saci_1116 with *p*NPA was normalized to "1" (4.2 U/mg) for lysates of expression cultures without 5'-UTR, and other activities were normalized accordingly. **C**) β-Galactosidase LacS-CtSS (*Saccharolobus solfataricus*) production with different 5'-UTRs. Specific activity of LacS with *p*NPG in crude extracts of expression cultures lacking 5'-UTR was set to "1" (1.4 U/mg), and other activities were normalized correspondingly.

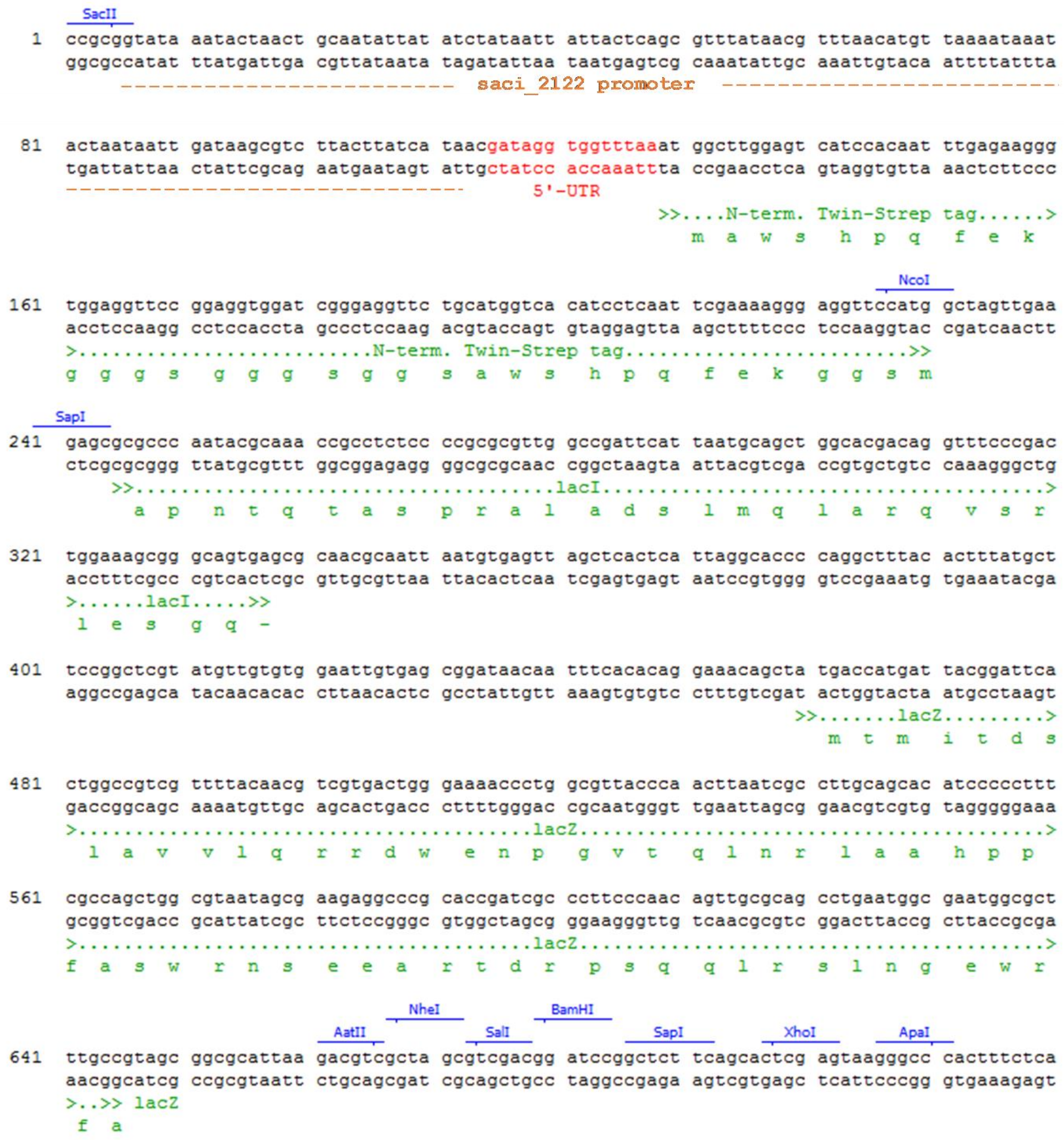


**Figure S8**: Important features of the pBSaraFX-*alba*UTR-NtSS expression vector. The following sequence features are indicated: the *saci_2122* promoter sequence, the inserted 5'-UTR sequence from *alba*, the nucleotide as well as derived amino acid sequence of the N-terminal located Twin-Strep tag, the *lacI* and *lacZ* fragments enabling blue-white screening in *E. coli* and the downstream multiple cloning site. Restriction sites are displayed. Clone Manager 7 (Sci Ed Software, USA) was used to generate the figure.

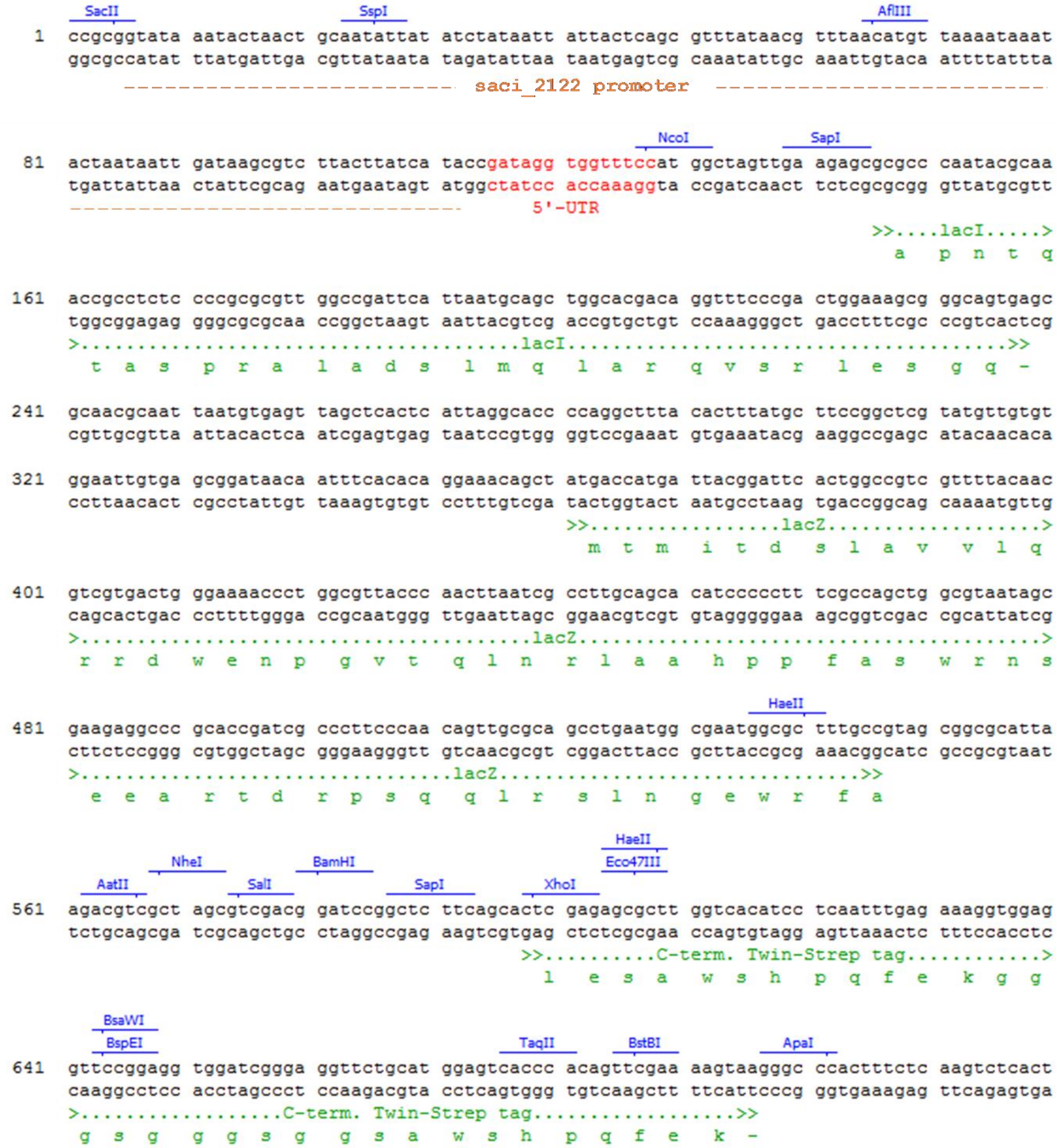


**Figure S9**: Important features of the pBSaraFX-*alba*UTR-CtSS expression vector. The following sequence features are indicated: the *saci_2122* promoter sequence, the inserted 5'-UTR sequence from *alba*, the *lacI* and *lacZ* fragments enabling blue-white screening in *E. coli*, the nucleotide as well as derived amino acid sequence of the C-terminal located Twin-Strep tag and the downstream multiple cloning site. Restriction sites are displayed. Clone Manager 7 (Sci Ed Software, USA) was used to generate the figure.

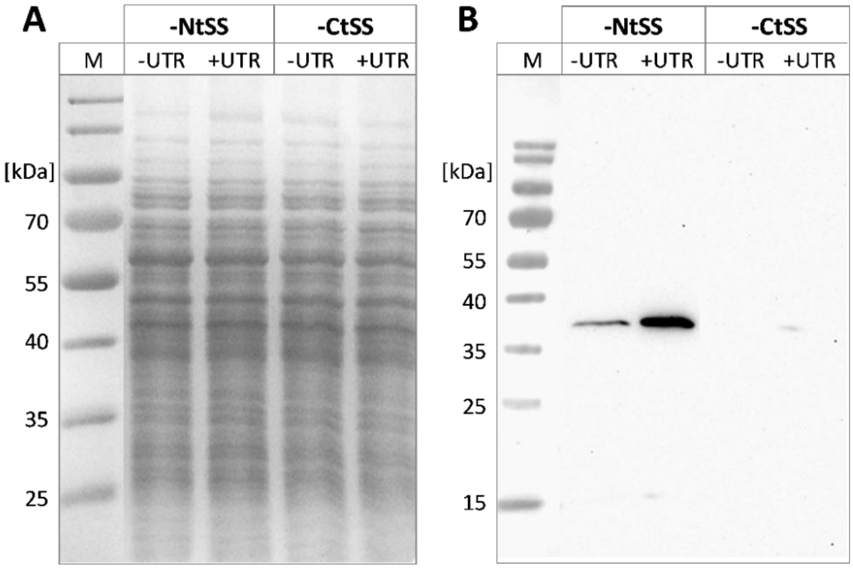


**Figure S10**: Effect of the *alba* 5'-UTR on the homologous production of glycosyltransferase Saci_1915. Protein production was compared without and with the use of *alba* 5'-UTR and with an N- or C-terminal Twin-Strep tag (-Nt/-CtSS). Expression plasmids pSVAaraFX-*saci_1915*-Nt/CtSS (**-UTR**) and pBSaraFX-*alba*UTR-*saci_1915*-Nt/CtSS (**+UTR**) were used. **A**) Coomassie-stained SDS-PAGE gel of whole cell samples. For all cell samples, an $OD_{600nm}$ value of 10 was adjusted and 15 µL of each sample were applied to SDS-PAGE. **B)** Immunodetection of Twin-Strep-tagged Saci_1915 using Strep-Tactin-HRP conjugate. Cultivation was performed as described in the legend of Fig. S3. Theoretical molecular weights (MWs): Saci_1915-NtSS: 40.8 kDa, Saci_1915-CtSS: 40.6 kDa. M: marker (protein ladder).

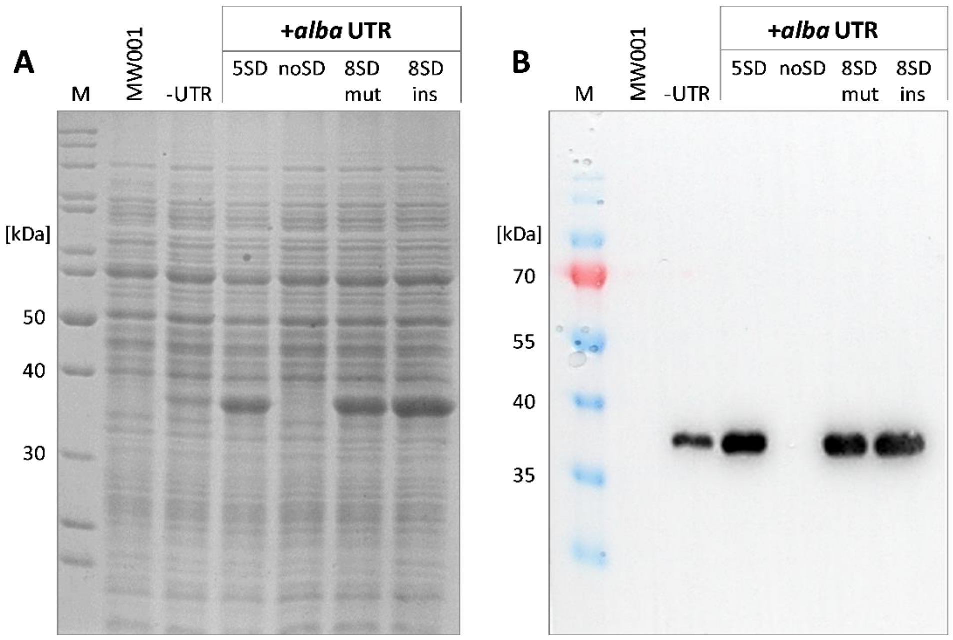


**Figure S11**: Influence of SD motif variation within *alba* 5'-UTR on esterase production. **A**) Protein visualization in cell lysates by SDS-PAGE and Coomassie staining. **B**) Immunodetection of C-terminal tagged esterase in cell lysate samples (Strep-Tactin HRP conjugate). Saci_1116-CtSS production was conducted using constructs without (-UTR) or with different variants of *alba* 5'-UTR (***+alba* UTR**), with modifications to the 5'-UTR sequence regarding the SD motif (Fig. 6, main text). M: marker (protein ladder).

**Table S1**: Primers used in this study. 5'-UTR sequences are underlined. Relevant nucleotides for the *alba* 5'-UTR modification by QuikChange PCRs® are indicated by the use of bold letters.

| Name | Sequence (5' → 3') | Description |
|---|---|---|
| **Insertion of 5'-UTR sequences into pSVAaraFX-*saci_1116*-CtSS** | | |
| UTR-fwd | CGTATTACCGCCTTTGAGTG | Fwd-primer for the insertion of 5'-UTRs |
| *saci_1322*-UTR-rev | ATACCATGGAAACCACCTATCGGTATGATAAG TAAGACGCTTATC | Insertion of *saci_1322* (Alba) 5'-UTR, *Nco*I site |
| *saci_0330*-UTR-rev | ATACCATGGTTGGGTCACCTTTTGGTATGATAAG TAAGACGCTTATC | Insertion of *saci_0330* 5'-UTR, *Nco*I site |
| *saci_1401*-UTR-rev | ATACCATGGGAGACACCGTTTCTGGTATGATAAG TAAGACGCTTATC | Insertion of *saci_1401* (Thα) 5'-UTR, *Nco*I site |
| *saci_2354*-UTR-rev | ATACCATGGACATACACCCGATATGTTATAA ATTGGTATGATAAGTAAGACGCTTATC | Insertion of *saci_2354* (SlaB) intergenic region, *Nco*I site |
| *saci_2355*-UTR-rev | ATACCATGGTTTTCACTTTCGGTATGATAAG TAAGACGCTTATC | Insertion of *saci_2355* (SlaA) 5'-UTR, *Nco*I site |
| **QuikChange PCR® primers for modification of *alba*UTR in pBSaraFX-*alba*UTR-*saci_1116*-CtSS** | | |
| QC-*alba*UTR-AA-fwd | GATAGGTGGTTT**AA**ATGGCTCCTTTAGATCCAACCA TAAAG | Modification of -2 and -1 nucleotides to "AA" |
| QC-*alba*UTR-AA-rev | GGAGCCAT**TT**AAACCACCTATCGGTATGATAAGTAA GACG | |
| QC-*alba*UTR-AC-fwd | GATAGGTGGTTT**AC**ATGGCTCCTTTAGATCCAACCA TAAAG | Modification of -2 and -1 nucleotides to "AC" |
| QC-*alba*UTR-AC-rev | GGAGCCAT**GT**AAACCACCTATCGGTATGATAAGTAA GACG | |
| QC-noSD-fwd | GCGTCTTACTTATCATACCGAT**TCCAC**GTTTCCATGGCTCCTT TAGATC | Complete mutation of SD motif (5 nts) |
| QC-noSD-rev | GATCTAAAGGAGCCATGGAAAC**GTGGA**ATCGGTATGATAAGT AAGACGC | |
| QC-8SDmut-fwd | GCGTCTTACTTATCATACCG**TG**AGGTG**A**TTTCCATGGCTCCT TTAGATC | Mutation of *alba* 5'-UTR nucleotides for establishment of 8 nt SD motif |
| QC-8SDmut-rev | GATCTAAAGGAGCCATGGAAA**T**CACCT**CA**CGGTATGATAAGT AAGACGC | |

| Name | Sequence (5' → 3') | Description |
|---|---|---|
| **Generation of pBSaraFX-*alba*UTR-8SDins-*saci_1116*-CtSS**<br>(8nt long SD motif within *alba* 5'-UTR by nucleotide insertions) | | |
| SD-fwd | ATGTTCTTTCCTGCGTTATC | Insertion of modified *alba* 5'-UTR (8SDins) in pSVAaraFX-*saci_1116*-CtSS |
| 8SD-ins-rev | ATACCATGGAAAC**T**CACCT**C**ATCGGTATGATAAGTAAGACGCTTATC | |
| **Generation of cloning vector pBSaraFX-*alba*UTR-NtSS** | | |
| *saci_1322*-UTR-NtSS-fwd | GATAGGTGGTTTAAATGGCTTGGAGTCATCCACAATTTG | Fwd-primer for insertion of *saci_1322* (*alba*) 5'-UTR in pSVAaraFX-NtSS (overlap PCR) |
| *saci_1322*-UTR-NtSS-rev | AGCCATTTAAACCACCTATCGTTATGATAAGTAAGACGCTTATC | Rev-primer for insertion of *saci_1322* (*alba*) 5'-UTR in pSVAaraFX-NtSS (overlap PCR) |
| **Amplification of genes of interest** | | |
| *Saci_1116*-*Nco*I-fwd | TAACCATGGCTCCTTTAGATCCAACCATAAAG | Amplification of *saci_1116*, addition of *Nco*I and *Xho*I restriction sites |
| *Saci_1116*-*Xho*I-rev | TATCTCGAGAAAACTGTTGTAGAATACTCGTCTC | |
| *lacS*-SSO-*Nco*I-fwd | GCGCCATGGACTCATTTCCAAATAGCTTTAG | Amplification of *lacS* from *S. solfataricus* for cloning of pBSaraFX-xxxUTR-*lacS*-CtSS |
| *lacS*-SSO-*Xho*I-rev | GTTCTCGAGGTGCCTTAATGGCTTTAC | |
| *Saci_1911*-*Nco*I-fwd | TATCCATGGTGTGGTCGATTGAGATCCC | Amplification of *saci_1911*, addition of *Nco*I and *Xho*I restriction sites |
| *Saci_1911*-*Xho*I-rev | CGGCTCGAGTCCTAATAAGTAGCCTAATATG | |
| *Saci_1915*-*Nco*I-fwd | TATCCATGGTGCCCAAAGACTTCAGTG | Amplification of *saci_1915*, addition of *Nco*I and *Xho*I restriction sites |
| *Saci_1915*-*Xho*I-rev | GCGCTCGAGACTAAACATCGAATTACTCC | |

**Table S2**: Effect of *alba* 5'-UTR insertion as well as changes in -2 and -1 nucleotide identities on esterase reporter protein (Saci_1116-CtSS) amounts. To determine the share of Saci_1116 in the total cell protein of *S. acidocaldarius* [%] the determined specific activities of Saci_1116-CtSS in *S. acidocaldarius* cell lysate samples (compare Fig. 4 in the main text) were set into relation to the enzyme's specific activity (124.5 U/mg). Fold changes are calculated as mean values, comparing the amount of reporter protein in lysate samples without any 5'-UTR to the different *alba* 5'-UTR variants (i.e. -2 and -1 nucleotides: AA, AC, CC). Expression cultures were harvested at $OD_{600nm}$ values of 0.6 (log growth phase).

| **Nucleotide identity at -2 and -1 positions of *alba* 5'-UTR** | **Specific activity in crude lysates [U/mg]** | **Amount of Saci_1116 in total protein [%]** | **Mean value fold change (-/+UTR)** |
|---|---|---|---|
| no UTR | 3.4 ± 0.2 | 2.6 ± 0.1 | - |
| AA | 16.0 ± 2.3 | 12.7 ± 1.8 | 4.7 |
| AC | 14.4 ± 1.1 | 11.5 ± 0.9 | 4.2 |
| CC | 13.1 ± 0.8 | 10.4 ± 0.7 | 3.9 |

**Table S3**: Student's t-test to assess the significant difference in reporter protein activity in dependency of the absence or presence of different *alba* 5'-UTR sequence variations. Significance was assessed for the data shown in Fig. 4B. The nucleotide identities of -2 and -1 nucleotides, located directly upstream of the translation start codon, are stated. The null hypotheses were stated as follows: i) the esterase activity in cell lysate samples is the same in the absence (no UTR) or presence of the *alba* 5'-UTR in the expression cassette; ii) the esterase activity is the same for different *alba* 5'-UTR sequence variants (different -2 and -1 nucleotide identities). The null hypotheses were rejected at 95% confidence level when the p-value was smaller than 0.05, indicating a statistically significant difference between the two conditions compared (indicated with an asterisk).

| **UTR 1** | **UTR 2** | **Mean specific activity with UTR 1 [U/mg]** | **Mean specific activity with UTR 2 [U/mg]** | **p-value** |
|---|---|---|---|---|
| no UTR | Alba UTR, AA | 3.4 | 16.0 | < 0.001 * |
| | Alba UTR, AC | | 14.4 | < 0.001 * |
| | Alba UTR, CC | | 13.1 | < 0.001 * |
| Alba UTR, AA | Alba UTR, AC | 16.0 | 14.4 | 0.34 |
| | Alba UTR, CC | | 13.1 | 0.10 |
| Alba UTR, AC | Alba UTR, CC | 14.4 | 13.1 | 0.16 |

**Table S4**: Selected UTR sequences from *S. acidocaldarius*. Information on the gene ID, the current annotation, the original UTR sequence in the genome of *S. acidocaldarius* as well as the adapted and inserted UTR sequence are given ("CC" at -2 and -1 positions (underlined) for maintenance of *Nco*I site in *saci_1116*-CtSS expression constructs). SD motifs are shown in bold.

| Gene | Protein | Genome UTR sequence | Inserted UTR sequence |
|---|---|---|---|
| *saci_0330* | Unknown function | AAA**AGGTGA**CCCAAAG | AAA**AGGTGA**CCCAACC |
| *saci_1401* | Thermosome subunit α (Thα) | AGAAAC**GGTG**TCTCAA | AGAAAC**GGTG**TCTCCC |
| *saci_2354* | S-layer protein B (SlaB) | AATTTATAACATATCG**GGTG**TATGTGT | AATTTATAACATATCG**GGTG**TATGTCC |
| *saci_2355* | S-layer protein A (SlaA) | GAAA**GTGA**AAAGT | GAAA**GTGA**AAACC |
| *saci_1322* | Alba | GAT**AGGTG**GTTTAA | GAT**AGGTG**GTTTCC |

**Table S5**: Effect of the different 5'-UTR sequences on the protein amounts of reporter protein Saci_1116-CtSS. To determine the share of Saci_1116 in the total cell protein of *S. acidocaldarius* [%] the determined specific activities of Saci_1116-CtSS in *S. acidocaldarius* cell lysate samples (compare Fig. 5 in the main text) were set into relation to the enzyme's specific activity (124.5 U/mg). Fold changes are calculated as mean values, comparing the amount of reporter enzyme in lysate samples without any 5'-UTR sequence to the amount of POI produced under the influence of a 5'-UTR. Expression cultures were harvested at $OD_{600nm}$ values of 0.8-1.2 (end of log growth phase).

| 5'-UTR | Specific activity in crude lysates [U/mg] | Amount of Saci_1116 in total protein [%] | Mean value fold change (-/+UTR) |
|---|---|---|---|
| no UTR | 4.2 ± 0.8 | 3.2 ± 0.7 | - |
| saci_0330 | 14.2 ± 3.1 | 11.3 ± 2.5 | 3.4 |
| Thα | 10.5 ± 1.4 | 8.4 ± 1.2 | 2.5 |
| SlaB | 10.1 ± 1.3 | 8.0 ± 1.0 | 2.4 |
| SlaA | 9.3 ± 1.2 | 7.4 ± 1.0 | 2.2 |
| Alba | 12.3 ± 1.1 | 9.8 ± 0.9 | 3.0 |

**Table S6**: Student's t-test to assess significant differences in reporter protein activity as a function of different 5'-UTR sequences. Significance was evaluated for the data shown in Fig. 5B. The null hypotheses were stated as follows: i) esterase activity in cell lysate samples is the same in the absence (no UTR) or presence of a 5'-UTR in the plasmid-based expression cassette; ii) the esterase activity is the same using different 5'-UTRs. The null hypotheses were rejected at 95% confidence level when the p-value was smaller than 0.05, indicating that there was a statistically significant difference between the two conditions compared (indicated with an asterisk).

| UTR 1 | UTR 2 | Mean specific activity with UTR 1 [U/mg] | Mean specific activity with UTR 2 [U/mg] | p-value |
|---|---|---|---|---|
| no UTR | saci_0330 | 4.2 | 14.2 | < 0.01 * |
| | Thα | | 10.5 | < 0.01 * |
| | SlaB | | 10.1 | < 0.01 * |
| | SlaA | | 9.3 | < 0.01 * |
| | Alba | | 12.3 | < 0.001 * |
| saci_0330 | Thα | 14.2 | 10.5 | 0.1 |
| | SlaB | | 10.1 | 0.1 |
| | SlaA | | 9.3 | 0.05 * |
| | Alba | | 12.3 | 0.3 |
| Thα | SlaB | 10.5 | 10.1 | 0.7 |
| | SlaA | | 9.3 | 0.3 |
| | Alba | | 12.3 | 0.2 |
| SlaB | SlaA | 10.1 | 9.3 | 0.45 |
| | Alba | | 12.3 | 0.1 |
| SlaA | Alba | 9.3 | 12.3 | 0.05 * |
